# Arterial vasodilation drives convective fluid flow in the brain: a poroelastic model

**DOI:** 10.1101/2021.09.23.461603

**Authors:** Ravi Teja Kedarasetti, Patrick J. Drew, Francesco Costanzo

## Abstract

The movement of fluid into, through, and out of the brain plays an important role in clearing metabolic waste. However, there is controversy regarding the mechanisms driving fluid movement, and whether the movement metabolic waste is primarily driven by diffusion or convection. The dilation of penetrating arterioles in the brain in response to increases in neural activity (neurovascular coupling) is an attractive candidate for driving fluid circulation, as it drives deformation of the brain tissue and of the paravascular space around arteries, resulting in fluid movement. We simulated the effects of vasodilation on fluid movement into and out of the brain using a novel poroelastic model of brain tissue. We found that arteriolar dilations could drive convective flow through the brain radially outward from the arteriole, and that this flow is sensitive to the dynamics of the dilation. Simulations of sleep-like conditions, with larger vasodilations and increased extracellular volume in the brain showed enhanced movement of fluid from the paravascular space into the brain. Our simulations suggest that both sensory-evoked and sleep-related arteriolar dilations can drive convective flow of cerebrospinal fluid from the paravascular space into the brain tissue around arterioles.

## Introduction

The circulation of cerebrospinal fluid (CSF) is thought to play the important role of clearing harmful solutes like amyloid-β from the brain (Bradbury, Cserr and Westrop, 1981; Cserr, Harling-Berg and Knopf, 1992; Weller, Kida and Zhang, 1992; Weller, 1998; Louveau *et al.*, 2015). The accumulation of these solutes in the brain extracellular spaces (ECS) has been linked to neurodegenerative diseases like Alzheimer’s (Hardy and Higgins, 1992; Selkoe and Hardy, 2016) and cerebral amyloid angiopathy (Yamada, 2000, 2015). The fluid-filled paravascular spaces (PVS) surrounding arteries and arterioles in the brain could provide a low resistance pathway for fluid and solute exchange between the CSF in the subarachnoid space (SAS) and the interstitial fluid (ISF) in the ECS, thereby playing a key role in the clearance of harmful metabolites. Studies in mice have shown that dyes injected into the cisterna magna or in the ventricles of the brain enter the ECS of the cerebral cortex primarily along the PVS of arterioles, suggesting that the PVS is the preferred pathway of solute exchange between CSF and ISF (Iliff *et al.*, 2012; Jeffrey J. Iliff *et al.*, 2013; Nedergaard, 2013). However, the nature and drivers of solute transport through the PVS remains controversial (Jin *et al.*, 2013; Asgari, De Zélicourt and Kurtcuoglu, 2016; Smith *et al.*, 2017; Holter *et al.*, 2017; Abbott *et al.*, 2018; Mestre, Tithof, *et al.*, 2018; Iliff and Simon, 2019; Smith and Verkman, 2019; Kedarasetti *et al.*, 2020; Kedarasetti, Drew and Costanzo, 2020; Rasmussen, Mestre and Nedergaard, 2021). While experimental data is key to understanding the fluid flow in the brain as it is direct physical evidence, there are certain limitations and artifacts to experimental methods (Mestre, Mori and Nedergaard, 2020). For example, with the currently available fluid tracing methods, fluid motion in the PVS and ECS can only be measured under Ketamine/Xylazine anesthesia but not in the awake state (Mestre, Tithof, *et al.*, 2018; Raghunandan *et al.*, 2021), and the insertion of glass pipettes and needles etc. into the brain parenchyma can appreciably alter the fluid flow in the brain (Mestre, Hablitz, *et al.*, 2018). In view of these limitations, mathematical modeling can be a valuable auxiliary tool in understanding the mechanisms driving fluid flow and solute transport through the PVS.

Several mathematical models have attempted to understand the nature and drivers of fluid and solute transport through the PVS (Bilston *et al.*, 2003; Schley *et al.*, 2006; Wang and Olbricht, 2011; Asgari, De Zélicourt and Kurtcuoglu, 2015, 2016; Holter *et al.*, 2017; Martinac and Bilston, 2019; Thomas, 2019; Daversin-Catty *et al.*, 2020; Kedarasetti *et al.*, 2020; Kedarasetti, Drew and Costanzo, 2020). However, the majority of published models of transport through the PVS have only simulated the fluid dynamics in the PVS in isolation (Bilston *et al.*, 2003; Wang and Olbricht, 2011; Asgari, De Zélicourt and Kurtcuoglu, 2016). These models only simulate fluid flow due to volume changes in the PVS that are directly driven by the movement the arteriolar walls (green region in Fig 1a). However, the effect of pressure changes in the PVS on the deformation of the surrounding ultrasoft brain tissue (Budday *et al.*, 2019) is almost never taken into account, thereby ignoring the feedback effect that volume and shape changes of the PVS has on the very geometry within which fluid flow occurs. In recent work (Kedarasetti *et al.*, 2020), we addressed this limitation by using fluid-structure interaction models to simulate the effect of brain elasticity on fluid exchange between the PVS and the SAS (pink region in Fig. 1a). The fluid-structure interaction models we used assumed that only one phase (fluid or solid) was present in the spatial domain said phase occupied. That is, the elastic response of the connective tissue in the fluid-filled spaces (SAS and PVS) and the fluid flow in the ECS were not simulated.

**Fig. 1:**
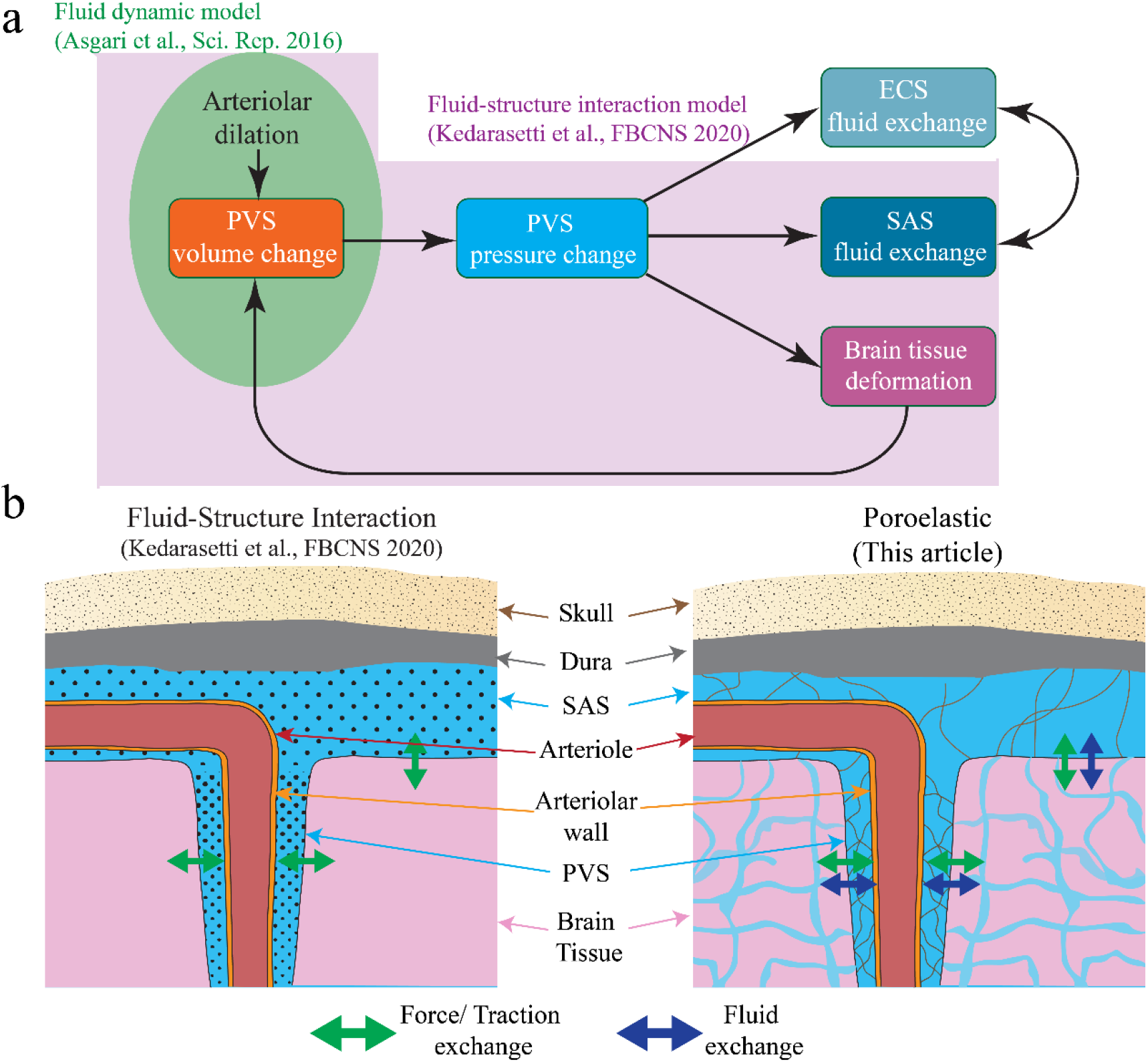
Schematic showing the working of a poroelastic model of the PVS, SAS and the brain tissue. **a.** Flowchart showing the full range of physics at play between the PVS, SAS, brain tissue, and the ECS that can be simulated by a poroelastic model. The field in green represents the physics that traditional fluid dynamic models capture (cf. Asgari et al., 2016). The field light purple (which contains the field in green) represents the physics captured by traditional fluid-structure interaction models (cf. Kedarasetti et al., 2020). The model presented in this paper extends the physics captured within the light purple field to also include the physics represented by the arrows outside said field. **b.** The advantages of using a poroelastic model over a traditional fluid-structure interaction model. In our previous fluid-structure interaction model we only simulated the fluid phase in the PVS and the SAS (shown by black dots). By contrast, with a poroelastic model we can also simulate the elasticity of the connective tissue and, more importantly, the fluid flow and transport *through* the ECS. These differences mean that a poroelastic model can simulate fluid exchange between the brain parenchyma and other fluid spaces *along with* the force exchange that can be simulated by a fluid-structure interaction model.

In this study, we improve on our previous modeling of transport through the PVS and brain by using 3D poroelastic models. Poroelastic models based on mixture theory (Bowen, 1976, 1980; Costanzo and Miller, 2017) can simultaneously simulate the solid and fluid phases in the same spatial domain, and the interactions between them. Using poroelastic models, simulations can be made of the volume changes of the PVS due to arteriolar wall movements and the resulting pressure changes in the PVS, which can drive fluid exchange between the PVS and the SAS or the ECS, and deform the brain tissue. Moreover, the poroelastic models can capture fluid exchange between the ECS and the SAS (Fig. 1a). Another way of thinking about the advantage of using a poroelastic model over a traditional fluid-structure interaction model is that while fluid-structure interaction models can only simulate the force transfer between the fluid filled regions (the SAS and the PVS) and the brain parenchyma, poroelastic models can additionally simulate the fluid-mass transfer between the fluid-filled regions and the brain parenchyma (Fig. 1b).

Using 3D poroelastic models, we considered two modes of solute transport through the PVS: dispersion and convection. Dispersion could improve solute transport over diffusion by oscillatory fluid exchange between the PVS and the SAS or the PVS and the ECS, while convection can drive directional fluid and solute transport from the SAS to the ECS, via the PVS. Several published models of transport through the PVS suggest that dispersion is the main mechanism of solute transport through the PVS (Asgari, De Zélicourt and Kurtcuoglu, 2016; Martinac and Bilston, 2019; Kedarasetti *et al.*, 2020). Dispersion-based solute transport is theoretically possible by any oscillatory movement of the arteriolar walls, like heartbeat-driven pulsations, intrinsic vasomotion of arteries (Winder *et al.*, 2017; Das, Murphy and Drew, 2021) and vasodilation due to increased neural activity, which have all been proposed as possible drivers of CSF flow (Bedussi *et al.*, 2017; Mestre, Tithof, *et al.*, 2018; von Holstein-Rathlou, Petersen and Nedergaard, 2018; van Veluw *et al.*, 2020). However, calculations based on fluid dynamics suggest that dispersion, with purely oscillatory flow would be a very ineffective means of solute transport in the PVS, while convection (even with low mean fluid velocities, of the order of 0.1 μm/s) would result in faster solute transport (Troyetsky *et al.*, 2021). Therefore, in this study, we focus on convective fluid flow from the PVS to the ECS and the possible drivers of this convective flow.

In this study, we demonstrate the possibility of convective transport through the PVS driven by a combination of the non-linear flow response of the fluid spaces in the brain and asymmetry in the waveform of arterial wall motions. The possibility of directional fluid flow through the PVS has been previously explored through mathematical models and numerical simulations (Bilston *et al.*, 2003; Schley *et al.*, 2006; Wang and Olbricht, 2011; Asgari, De Zélicourt and Kurtcuoglu, 2016; Daversin-Catty *et al.*, 2020; Kedarasetti, Drew and Costanzo, 2020). However, most of the published models only considered the peristaltic motion of arteries driven by heartbeat pulsations as the possible driver of convectional transport. While it is theoretically possible to drive directional CSF flow by peristaltic pumping (Bilston *et al.*, 2003; Wang and Olbricht, 2011), models using realistic dimensions and boundary conditions representing the anatomy of the PVS (Asgari, De Zélicourt and Kurtcuoglu, 2016; Daversin-Catty *et al.*, 2020; Kedarasetti, Drew and Costanzo, 2020) suggest that heartbeat-driven pulsations of arteries drive mostly oscillatory flow in the mouse brain with negligible directional fluid flow. To the best of our knowledge, this study is the first to consider an alternative mechanism to peristalsis for driving convective transport through the PVS.

The role of functional hyperemia in driving fluid and solute transport through the PVS will be the major focus of this study. Functional hyperemia (Iadecola, 2017) is the dilation of arteries and arterioles in the brain in regions of increased neural activity, potentially driven by a subset of neurons. Though it is often stated that functional hyperemia is required to match the brain’s energetic demands (Leithner and Royl, 2014), this is not the case, and the underlying physiological purpose of functional hyperemia is a mystery. The hypothesis that functional hyperemia drives PVS solute transport has received some attention recently, with support from both experiments (von Holstein-Rathlou, Petersen and Nedergaard, 2018; van Veluw *et al.*, 2020) and theoretical models (Kedarasetti *et al.*, 2020). Using our 3D poroelastic models, we tried to understand how different characteristics of functional hyperemia affect solute transport through the PVS. Our models showed that the temporal characteristics of functional hyperemia, which usually consist of a rapid dilation of arterioles (reaching the peak dilation within two seconds) followed by a slower return to resting diameter over several seconds (Silva, Koretsky and Duyn, 2007; Drew, Shih and Kleinfeld, 2011; Gao, Greene and Drew, 2015; He *et al.*, 2018) could drive convective fluid flow. The models also showed that hyperemia during sleep, which has arteriolar dilations several times larger than those during the awake state (Bergel *et al.*, 2018; Turner *et al.*, 2020a), combined with the increased extracellular volume in the brain (Xie *et al.*, 2013) could explain the larger solute transport in the brain parenchyma observed during sleep (Xie *et al.*, 2013; Hablitz *et al.*, 2019). The models also suggest that the low frequency oscillations in vessel dilation during neural activity and sleep play a major role in solute transport through the PVS.

## Model assumptions

The geometry of the model was created to represent the anatomy of a single penetrating arteriole in the mouse cortex, while keeping the shape relatively simple. The dimensions of the model geometry are shown in Fig 2a. The entire geometry had a size of (*x* × *y* × *z*) 80 μm × 200 μm × 200 μm, with the *z* direction being perpendicular to the pial surface. The model was composed of two domains, one representing the fluid-filled SAS and the PVS (translucent blue in Fig 2a) and the other representing the poroelastic brain tissue (pink in Fig 2a). To keep the geometry simple, the model simulated a segment of the brain from the cortical surface to 150 μm in depth (*z* direction), below which arterioles usually branch out into smaller arterioles or capillaries (Blinder *et al.*, 2013; Horton *et al.*, 2013; Gagnon *et al.*, 2015). The dimensions of the geometry in the *x* and *y* directions were chosen to represent half of the typical separation between arterioles in the cortex (Nishimura *et al.*, 2007; Gagnon *et al.*, 2015; Shih *et al.*, 2015; Adams *et al.*, 2018). The SAS of the model had a nominal width of 50 μm (Coles, Myburgh, *et al.*, 2017; Coles, Stewart-Hutchinson, *et al.*, 2017). The part of the geometry representing the SAS and the PVS was built with a cavity representing an arteriole penetrating into the brain. The segment of the arteriole passing through the SAS had a diameter of 20 μm (Drew *et al.*, 2010; Shih *et al.*, 2012), with its long axis along the *y*-axis of the model. The arteriole was assumed to penetrate into the brain tissue along the *z*-axis, with its diameter tapering down to 15 μm at 150 μm below the brain surface of the brain. The PVS surrounding the arteriole was assumed to be an annular region with a width of 8 μm near the surface of the brain and 5.5 μm at 150 μm below the brain surface. The dimensions of the PVS were taken from experimentally-determined values from published imaging data (Iliff *et al.*, 2012; Schain *et al.*, 2017; Mestre, Tithof, *et al.*, 2018). For the geometry of the PVS, a relatively simple annular shape was chosen instead of a more realistic eccentric and elliptical annular shape (Min Rivas *et al.*, 2020) to avoid further complicating the model by increasing the number of unknown parameters (like eccentricity), or by adding a cumbersome boundary condition at the common interface of the arteriole, the PVS, and the brain tissue. All the sharp corners in the model geometry were smoothened by using a circular fillet. The geometry was sliced in half at the *yz* plane (*x* = 0) to reduce the size of the calculations using symmetry boundary conditions (see the section on boundary conditions in Methods). The model was oriented so that the origin (0,0,0) was on the axis of the vessel and at the bottom surface of the brain parenchyma. A tetrahedral mesh was created for the half section with elements of thickness 2 μm at the surfaces representing the arteriolar wall, the skull and the interface between the fluid-filled spaces and the brain tissue. The mesh size was gradually increased to 10 μm (Fig. 2b).

**Fig 2:**
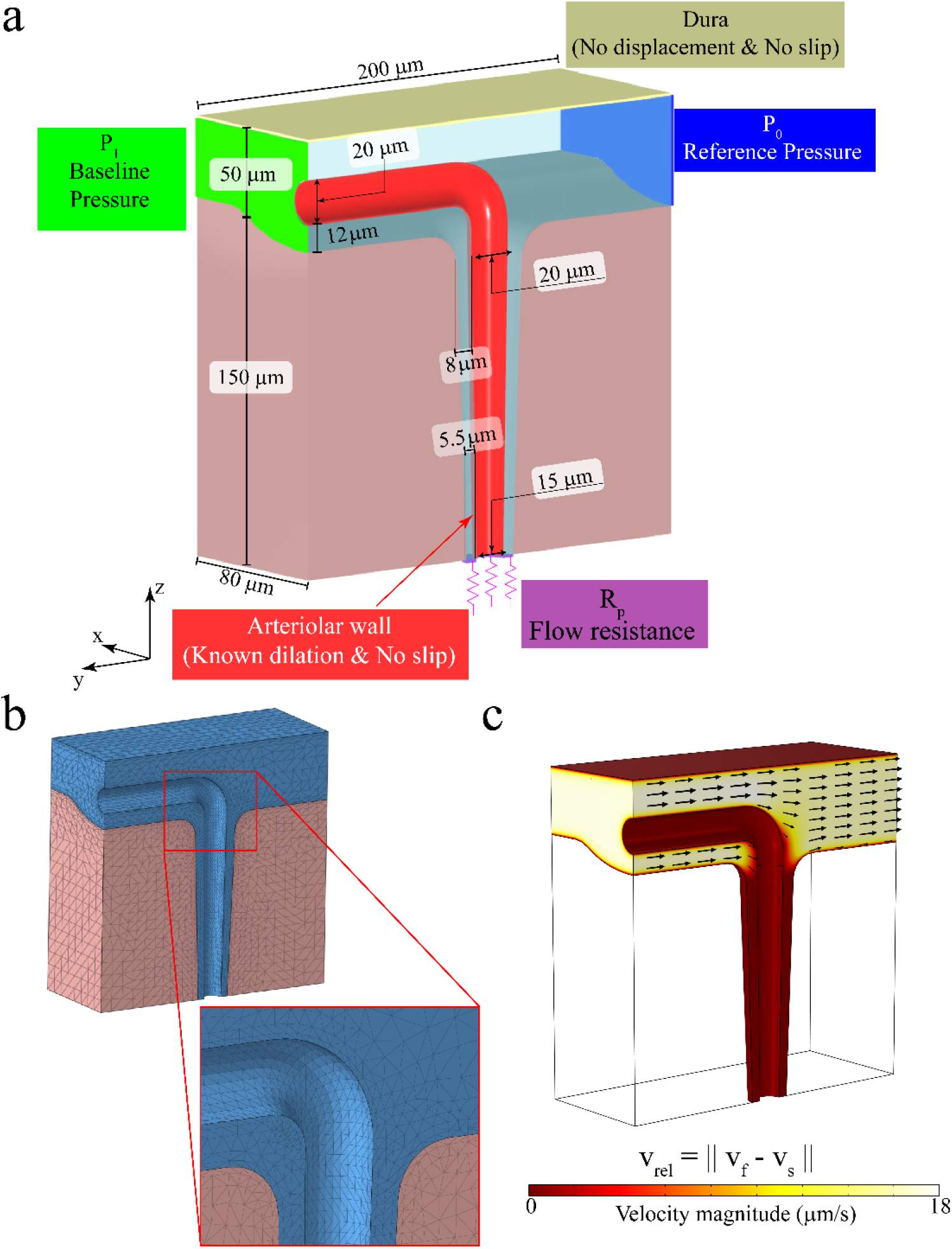
Geometry, boundary conditions and discretization of the model. **a.** The geometry of the model showing the two domains, with the dimensions and boundary conditions. Solid displacement and fluid velocity were prescribed at the red- and cream-colored surfaces. Pressure-like tractions were prescribed on the green- and blue-colored surfaces. Flow resistance (Robin) boundary co nditions were prescribed on the purple-colored surface. Symmetry boundary conditions were prescribed on all other surfaces. **b.** Tetrahedral mesh used for the finite element model. A fine mesh, with elements of 2 μm were used near the regions where no-slip boundary conditions were prescribed and at the interface between the two domains. The mesh size was gradually increased to 10 μm. **c.** The fluid flow in the SAS at the baseline state, which is a result of the pressure difference applied across the ends (green- and blue-colored surfaces in **(a)**).

The constitutive models and the model parameters were chosen to capture the experimentally determined mechanics of the brain tissue and the surrounding fluid spaces. We use the superscript ^1^ to represent the parameters in the fluid-filled spaces, and ^2^ to represent the parameters in the brain tissue. An incompressible Darcy-Brinkman model was used for fluid flow through porous spaces, with a mass density 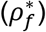 of 1000kg/m^3^ and viscosity (*μ_f_*) of 0.001Pa⋅s (Yetkin *et al.*, 2010; Støverud *et al.*, 2013). The fluid permeability, 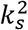, of the brain tissue was assumed to be 2 × 10^−15^ m^2^, based on experimental measurements (Neeves *et al.*, 2006; Smith and Humphrey, 2007). The permeability of the PVS, 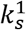 for z < 130 μm, was chosen to be 2 × 10^−14^ m^2^, 10 times higher than that of the ECS. The SAS in the model is supposed to represent a combination of the open (completely fluid-filled and not porous) PVS of surface arterioles (Mestre, Tithof, *et al.*, 2018; Min Rivas *et al.*, 2020) and the porous SAS, and therefore, a higher permeability, 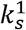 for z > 150 μm, of 2 × 10^−12^ m^2^ was used for the SAS. The permeability in the fluid-filled domain for 130 μm ≤ *z* ≤ 150 μm was transitioned using a function with continuous first and second derivatives (*step* function in COMSOL Multiphysics). The brain tissue fluid volume fraction 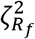 was set to 0.2 to represent the 20% of the brain volume occupied by the extracellular fluid, which is in the range of values measured by 2D and 3D electron microscopy data (Korogod, Petersen and Knott, 2015; Holter *et al.*, 2017), as well as measurements with real-time Iontophoresis (Xie *et al.*, 2013). For the fluid-filled spaces, a higher fluid volume fraction 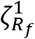 of 0.8 was used. An incompressible Neo-Hookean model was used for the solid phase. A shear modulus 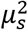 of 2kPa (Budday *et al.*, 2017; Mihai *et al.*, 2017; Weickenmeier *et al.*, 2018) was used for the brain tissue and a small shear modulus 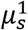 of 20Pa was used in the fluid-filled domain to represent the connective tissue in the fluid-filled spaces. A density of 1000kg/m^3^ was used for the solid phase (Barber, Brockway and Higgins, 1970).

The boundary conditions for the model were chosen to represent a segment of the cerebral cortex in an active region of the brain, i.e., a section of the cortex containing a dilating arteriole surrounded by other arterioles dilating with similar dynamics. This choice of boundary conditions (depicted in Fig. 2a) is apt for simulating both sleep and awake states, where arteries withing a few square millimeter patch will dilate (Yu *et al.*, 2014, 2016; Turner *et al.*, 2020b) simultaneously. The arteriolar wall motion was simulated by the radially outward solid displacement at the surface of the cavity representing the penetrating arteriole (red surface in Fig. 2a), and no-slip boundary conditions were used for the fluid. A small pressure difference was applied across the ends of the SAS in the form of traction forces on the fluid phases (*P*_0_ = 0mmHg on the blue surface and *P*_1_ = 0.01mmHg on the green surface in Fig. 2a), to simulate the flow driven by the secretion of CSF (and possibly by arterial pulsations). The value of *P*_1_ was chosen to achieve a maximum flow velocity of 20 μm/s around arterioles on the surface of the brain (Bedussi *et al.*, 2017; Mestre, Tithof, *et al.*, 2018), when the arteriole is at the baseline diameter (Fig. 2c). The bottom surface of the PVS (*z* = 0) was assumed to be connected to the brain parenchyma and the PVS of smaller arterioles and therefore a flow resistance, with a value of 10 times the flow resistance of the PVS was used (purple surface in Fig. 2a). This resistance was assumed to represent the flow resistance of all the pathways that fluid can flow through before re-entering the SAS, and therefore zero pressure was assumed beyond the resistor. A zero-displacement condition for the solid phase and a no-slip boundary condition for the fluid phase were implemented on the top surface of the SAS (*z* = 200 μm), which represents the skull-fixed dura. On all other free surfaces of the model, the solid displacement and fluid flow perpendicular to the surface were set to zero. For the *yz*-plane (*x* = 0), this boundary condition reflects the symmetry assumption, while for the surfaces at x = 80 μm, y = −100 μm and y = 100 μm the boundary condition reflects the assumption that the domains represented in the model are surrounded by similar structures experiencing similar arteriolar dilation. At the bottom surface of the brain tissue (*z* = 0), the condition of no fluid flow perpendicular to the surface was deemed more apt than a flow resistance boundary condition, as the latter would set uniform, flow-dependent traction across the whole surface. At the interface between the two domains, mass and momentum continuity are maintained by special boundary conditions, usually referred to as jump conditions (see interface conditions in Methods).

## Results

### Functional hyperemia can drive directional fluid flow through the PVS

We first examined the possibility of convective solute transport through the PVS driven by functional hyperemia and the factors contributing to the convective solute transport. Specifically, we quantified the contribution of the waveform of functional hyperemia to directional fluid flow through the PVS. To do this, we compared two modes of vasodilation, temporally symmetric and temporally asymmetric vasodilations. For symmetric vasodilation, the temporal waveform of arteriolar wall displacement resembles a Gaussian pulse, and the negative radial velocity of the arteriolar wall during the contraction of the vessel is equal and opposite to the positive radial velocity during dilation (Fig. 3a top). For the case of asymmetric dilation, the waveform of arteriolar wall displacement resembles that seen in functional hyperemia (Drew, 2019), with a sharper increase of vessel diameter at the beginning of the event, followed by a slow return to baseline (Fig. 3a bottom). In this case, the peak magnitude of the negative radial velocity of the arteriolar wall is roughly half the value of peak positive radial velocity. To quantify the convective flow driven by arteriolar dilation, we defined two time-averaged Peclet numbers (see Methods, Peclet numbers), averaged over 10s of simulation. The *axial* Peclet number, *Pe_a_*, was defined based on the time-averaged relative fluid velocity (i.e., relative to the solid) through the bottom face of the PVS, and represents directional pumping by arteriolar wall motions in the traditional sense, similar to peristaltic pumping. The *radial* Peclet number, *Pe_r_*, was defined based on the time-averaged radial component of the relative fluid velocity at the interface of the PVS and the brain tissue, and represents directional fluid flow into the ECS due to asymmetries in the flow resistances of the SAS-PVS-ECS system. The Peclet numbers were defined based on the diffusion coefficient of amyloid-β (*D_aβ_*, see Table 1) and a characteristic length of 150 μm.

**Fig. 3:**
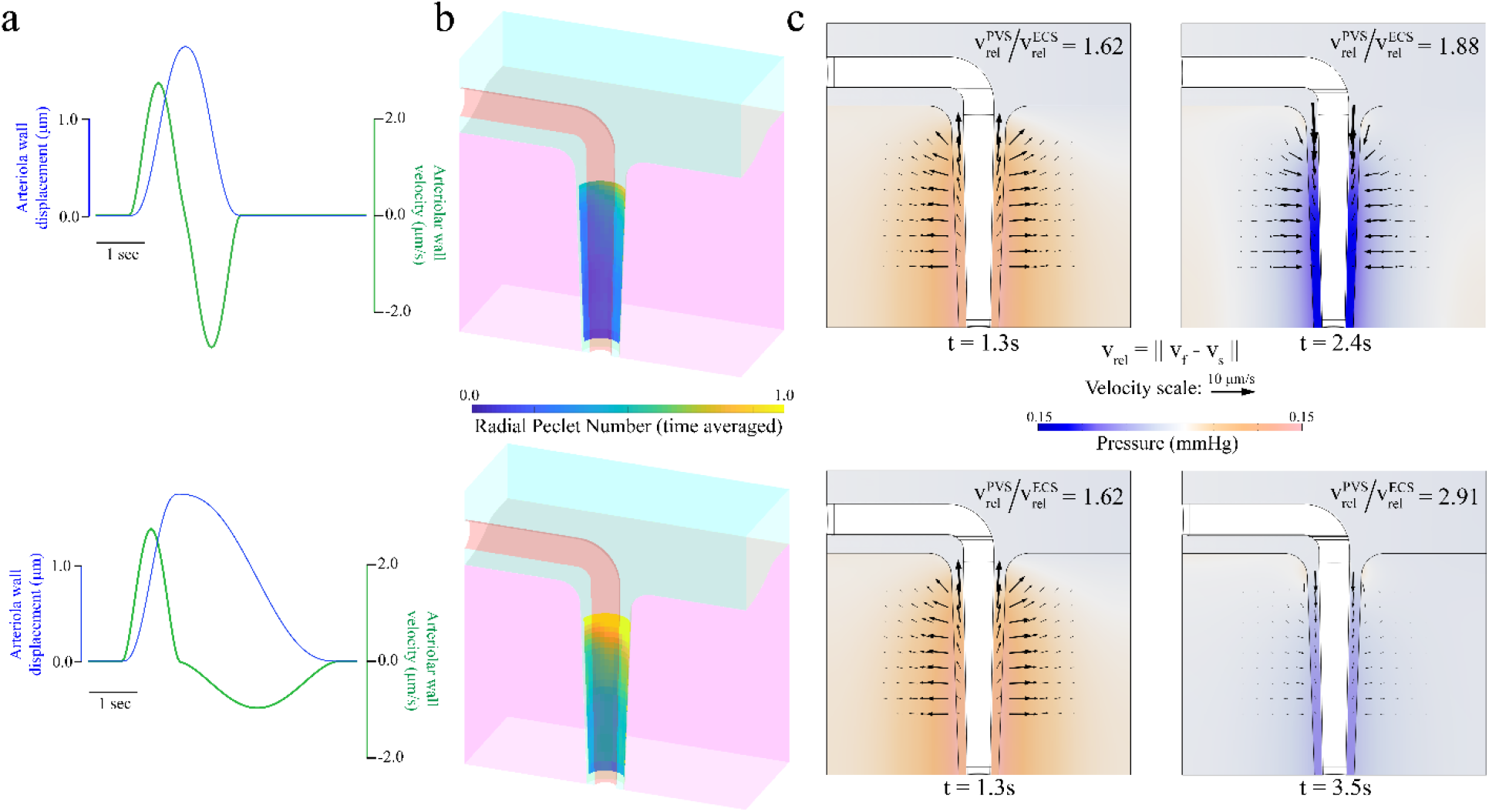
The asymmetric waveform of functional hyperemia can drive net directional fluid flow through the PVS. a. The radially outward displacement (blue) and velocity (green) of the arteriolar wall for the case of symmetric dilation (top) and asymmetric dilation (bottom). b. The time averaged radial Peclet numbers at the PVS-ECS interface as a result of symmetric (top) and (asymmetric) vasodilation. c. The pressure and relative fluid velocity in the PVS and the ECS at the times of maximum radially outward and inward arteriolar wall velocity for symmetric (top) and asymmetric (bottom) dilation. The colors show the pressure value in mmHg and the arrows show the magnitude and direction of relative fluid flow. By comparing the ratio of the maximum relative velocity in the PVS and SAS, it can be seen with asymmetric vasodilation more fluid enters the ECS through the PVS than returns into the PVS through the ECS.

**Table 1.**
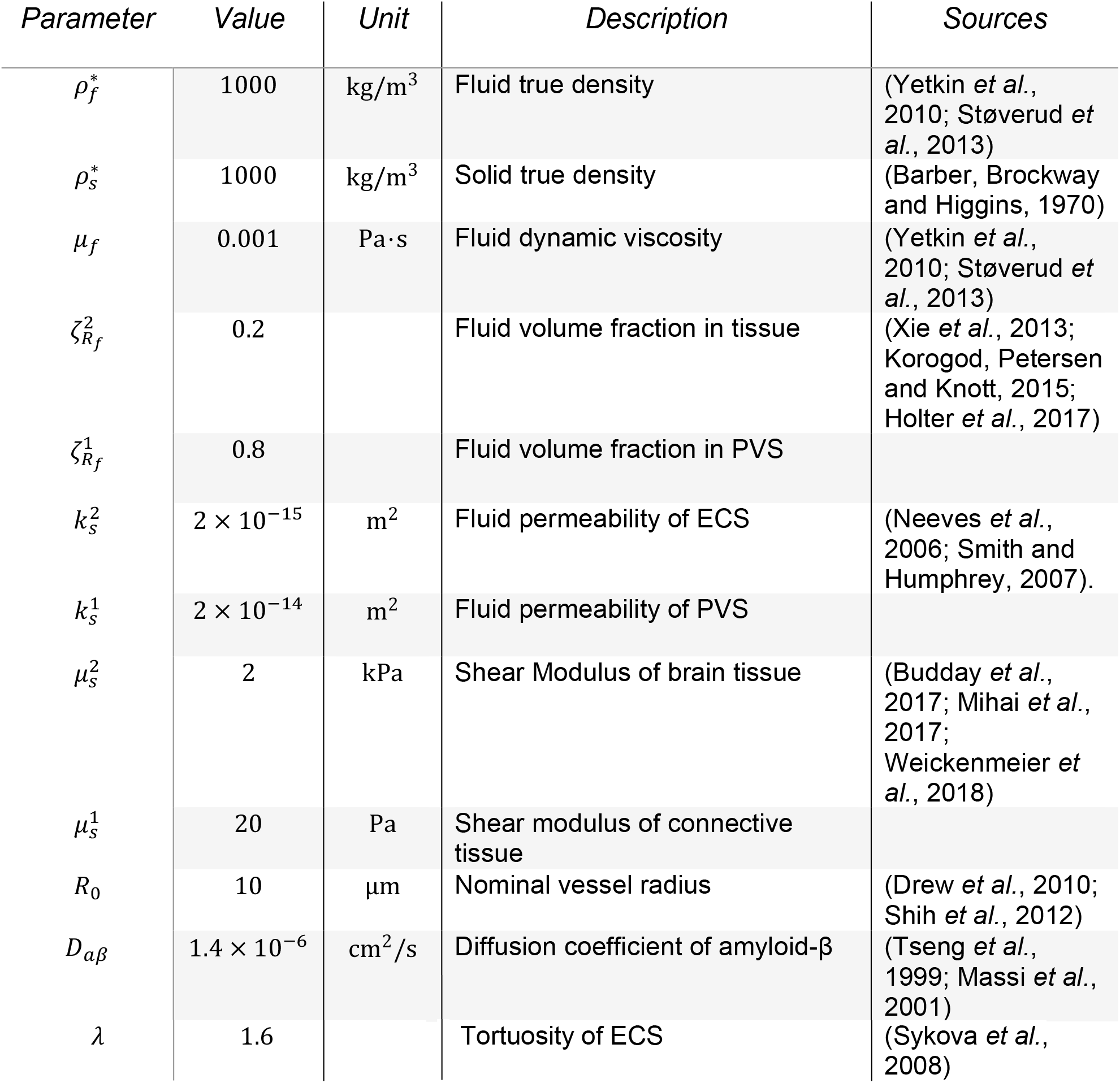
Model Parameters

Arteriolar dilation with an asymmetric waveform resulted in appreciable radially outward fluid flow into the ECS through the PVS, while neither waveform resulted in directional fluid flow in the axial direction. The time-averaged radial Peclet numbers at the interface between the brain tissue and the PVS are shown in Fig. 3b. For the symmetric dilation waveform, the maximum time-averaged radial Peclet number was 1.11 (Fig. 3b, top), while for the asymmetric waveform, the maximum time-averaged radial Peclet number was 2.07 (Fig. 3b, top). The time-averaged axial Peclet number for the symmetric waveform was 0.0003 and asymmetric waveform was 0.0012, indicating that arteriolar dilations cannot drive directional fluid flow through the PVS alone. This might be a result of using a standing-wave dilation in our simulations rather than a traveling-wave dilation, although previously published mathematical models that used realistic dimensions of the PVS (Asgari, De Zélicourt and Kurtcuoglu, 2016; Daversin-Catty *et al.*, 2020; Kedarasetti, Drew and Costanzo, 2020) suggest that traveling-wave dilations cannot drive directional fluid flow through the PVS alone.

The reason for more pronounced radially outward pumping from the asymmetric dilation compared to symmetric dilation is a result of the relative fluid velocity in the ECS during arteriolar dilation and contraction, and the ratio of the relative fluid velocities in the PVS to those in the ECS (Fig. 3c). In Fig. 3c, the arrows show the direction and magnitude of the relative fluid velocity in both the domains at the times of peak outward and inward velocity of the arteriolar wall. The ratio of the maximum velocity magnitudes (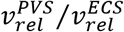 in Fig. 3c) in the PVS and ECS indicate the ratio of fluid in the PVS being exchanged with the SAS and the ECS respectively. For the case of symmetric dilation, this ratio of maximum velocity magnitudes is similar during dilation and contraction of the vessel, indicating that roughly the same amount of fluid leaving the PVS through the ECS returns into the PVS through the ECS. For the case of asymmetric dilation, the ratio of maximum velocity during contraction is nearly twice the ratio during dilation, indicating that during the slow contraction of the vessel, only a fraction of the fluid leaving the PVS through the ECS returns through the same path. Therefore, for each dilation and contraction there is a larger net directional flow from into the ECS through the PVS for the asymmetric dilation waveform compared to the symmetric dilation.

Note that the arrows plotted in Fig. 3c showing the relative fluid velocity are an indication of how far fluid travels in the PVS and the ECS, but not an indication of the amount of fluid entering or exiting the SAS. The amount of fluid moving can be better understood by examining the filtration velocities (Fig. S2). Filtration velocity is the relative fluid velocity multiplied the fluid volume fraction (See Eq. in Methods), and is an indicator of the amount of fluid flow relative to the solid phase. The conservation of fluid-mass dictates that the fluid flowing through the interface between the ECS and the PVS needs to be conserved, which is why the component of filtration velocity perpendicular to the interface between the two domains remains continuous in Fig. S2. The lower fluid volume fraction in the ECS means that for the same flowrate to be maintained, the fluid velocity in the ECS needs to be higher than the fluid velocity in the PVS. This is reflected in the higher magnitude relative fluid velocity in the ECS compared to the relative fluid velocity in the PVS, in Fig. 3c.

The fluid velocities in the PVS and the ECS seen in the model are a result of the difference in the response of poroelastic mixtures to volume and pressure changes. When an incompressible poroelastic mixture is subject to transient volume changes, the fluid flow response is approximately linear with the volume changes, and the fluid flow rate closely follows the volume changes. However, when the mixture is subject to transient pressure changes, the fluid flow response is highly non-linear with respect to the pressure, and the fluid flow rate changes “lag” behind the pressure changes. To demonstrate this phenomenon, we created a simple 2D poroelastic model (Fig. S1) of a square block of length 150 μm. The top and bottom edges of the square are subject to zero solid displacement in the vertical direction and no-slip boundary conditions for the fluid. At the right end, the horizontal solid displacement is set to zero, while no traction is applied on the fluid phase. When a Gaussian pulse of displacement (along with no-slip boundary condition) is applied at the left end, the fluid flowrate at the right end follows the applied wall velocity (derivative of the displacement) waveform. In contrast, when a pressure-like traction is applied on the left end (on both solid and liquid phases), there is a clear lag between the fluid flowrate at the right end the pressure waveform. This kind of lag between applied pressure and the flow response of poroelastic solids has been observed in soils (Senjuntichai and Rajapakse, 1993; Xie, Liu and Shi, 2004). When the arteriole dilates in our model, the fluid-filled domain is subject to volume changes, while the domain representing the parenchymal tissue is subject to pressure changes due to the volume changes in the fluid-filled domain. Therefore, while the PVS fluid outflow during arteriolar dilation occurs through both the SAS and the ECS, the PVS fluid inflow during the arteriolar contraction that follows dilation occurs more from the SAS, because the flow response through the ECS is lagging. This difference between the inflow and outflow pathways for PVS fluid is further enhanced with the asymmetric waveform, because the faster dilation drives larger pressure changes in the PVS, compared to the slower contraction that follows dilation.

The brain tissue deforms in the poroelastic model due to pressure changes in the PVS (Fig. S3). This deformation of the brain tissue was also predicted by our fluid-structure interaction model and demonstrated by our in-vivo imaging data (Kedarasetti *et al.*, 2020). The radially outward displacement of the brain tissue relative to the displacement of the arteriolar wall in the 3D poroelastic model (Fig. S3b-c) was smaller than the displacement predicted by the fluid-structure interaction model. There are two main reasons for this. First, the pressure in the PVS in the poroelastic model acts on both fluid and solid phases and drives fluid flow, unlike the fluid-structure interaction model where all the pressure is driving the deformation of the brain tissue. Second, the width of the PVS in this poroelastic model was in the higher range of possible values, while in our fluid-structure interaction model (Kedarasetti *et al.*, 2020) the width of the PVS was in the lower range of possible values, and therefore, because of the larger cross-sectional area of the PVS, for the same volume of fluid displaced by the arteriolar dilation, the fluid velocities and the resulting pressure changes in this poroelastic model are smaller than those in the fluid-structure interaction model. In the poroelastic model, there was also a displacement of the brain tissue in the z-direction (towards the surface) during arteriolar dilation (Fig. S3d-e). The displacement in the vertical direction was because of an “expansion” of the brain tissue when fluid from the PVS entered the ECS.

### Functional hyperemia can drive solute penetration into the brain

A common method for experimentally visualizing fluid movement into the brain is to inject tracers (either a fluorescent dye or particles) into the “large” CSF chambers in the cranial space (cisterna magna or the ventricles) and observe their movement (Iliff *et al.*, 2012; Ma *et al.*, 2017; Mestre, Hablitz, *et al.*, 2018; von Holstein-Rathlou, Petersen and Nedergaard, 2018). To connect the results of the simulations to experimental observations we modeled the fluid movement driven by arteriolar dilation by adding fluid particle tracking to the poroelastic simulations. It is important to keep in mind that there is a key difference between the movement of the particles that we are simulating here and the movement of physical tracers. These simulated particles are merely, passive tracers and do not have physical properties of their own. The simulated particles do not diffuse and have the same mobility as water, irrespective of the fluid volume and tortuosity changes in the PVS and the brain, unlike real tracers. The size of the particles in the figures do not correspond to the “real” size of the particles, they are for visualization purposes only. Diffusion equations were not added to the model to prevent further complicating the model. The physics at play is purely that of the computed fluid flow in the poroelastic mixture.

We started the particle tracking simulations with 243 equally spaced fluid particles (27 rings of 9 particles) in the PVS (Fig. 4a). The fluid particle motion was simulated for models with either symmetric or asymmetric vasodilation (Fig. 4b). These simulations had a duration of 60 seconds, where one vasodilation event occurred once every 10 seconds. At any given time in the simulation, the fluid particles were classified to be in the PVS, ECS or SAS based on their position. Area plots showing this distribution of particles (Fig. 4c), and 3D lines showing the particle trajectories for 60 seconds were plotted (Fig. 4d) to visualize and understand the physics of fluid motion through the fluid spaces surrounding a dilating arteriole.

**Fig. 4:**
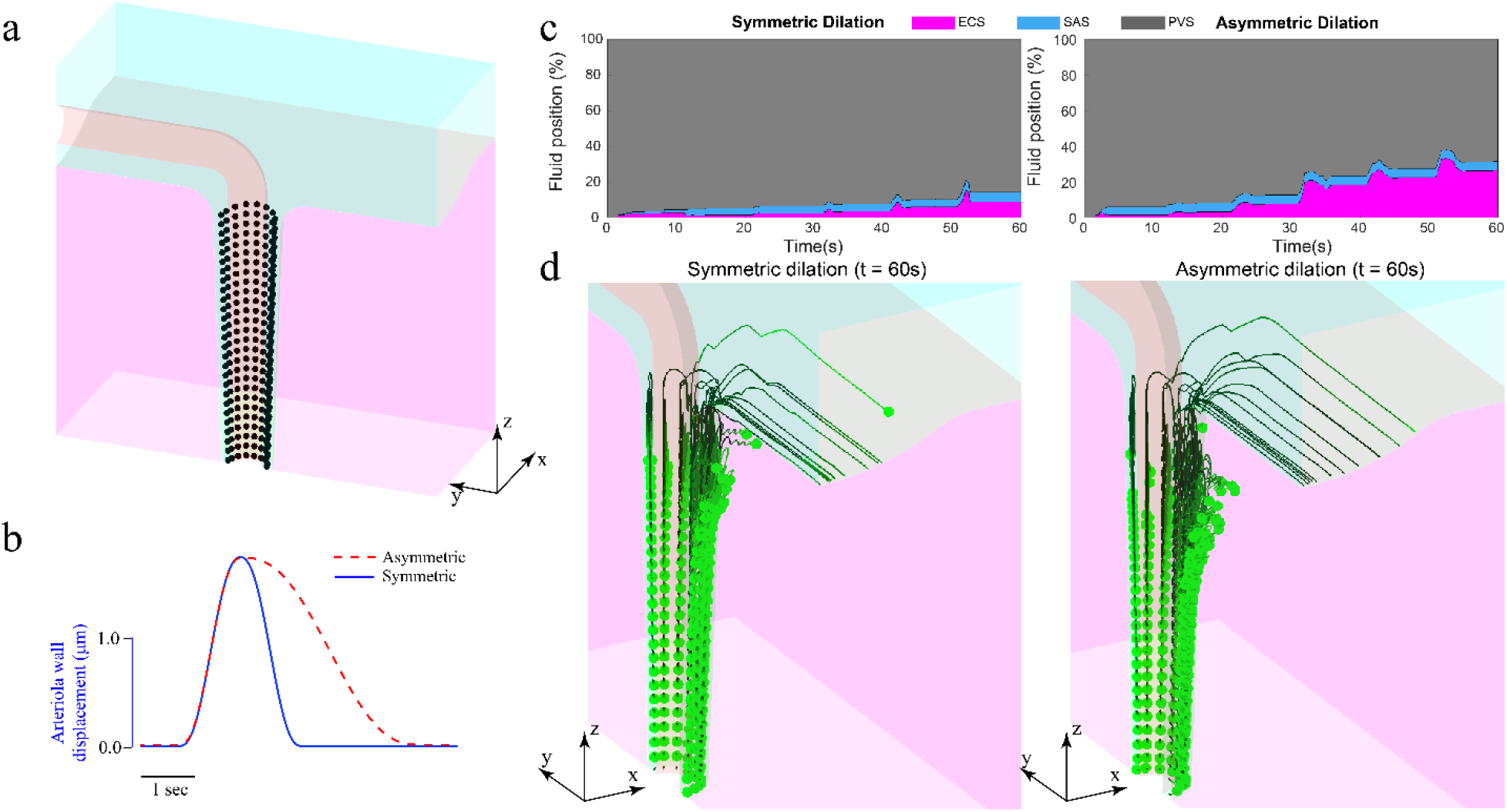
Functional hyperemia drives solute penetration into the brain. **a.** The initial position of particles used for particle tracking simulations. **b.** The waveform of radial displacement of arteriolar displacement for symmetric and asymmetric dilation in the model. The asymmetric dilation waveform r epresents functional hyperemia. **c.** The distribution of fluid position for 60 seconds of simulation with symmetric (left) and asymmetric (right) dilation waveform. Asymmetric dilation drives nearly three times (27%) PVS fluid movement into the brain compared to asymmetric dilation (9%). Both symmetric and asymmetric dilation drive similar PVS fluid movement into the SAS (5%). **d.** The particle trajectories of fluid particles shown in (a). Asymmetric dilation moves PVS fluid deeper below the surface into the EC S and moves the brain further into the brain.

The particle tracking simulations showed that the asymmetric waveform of functional hyperemia can drive appreciable solute penetration into the ECS, with nearly three times (26.75%) the fluid particles moving from the PVS to the ECS compared to vasodilation with a symmetric waveform (9%) (Fig. 4c) over the same amount of time. The models suggest that for a symmetric waveform, only PVS fluid close to the surface of the brain penetrates into ECS, while for the asymmetric waveform the PVS fluid deeper in the brain penetrates into the ECS (Fig. 4d). Moreover, the radial distance to which the dye penetrates into the brain is larger for asymmetric dilation compared to symmetric dilation. The particle tracking simulations show that the PVS fluid that moves into the SAS is the same for symmetric and asymmetric dilation, which is expected, as the fluid velocity during dilation is same for both cases (Fig. 3c left). Note that while the net flow of fluid is from the PVS into the ECS, there are times where the flow reverses.

The PVS fluid penetration into the ECS is not an artifact of the directional fluid flow imposed through the pressure difference across the two ends of the SAS (Fig. 2a). To verify this, we repeated the particle simulations with models where the pressure difference imposed across the ends of the SAS was 1/10^th^ of the value used in the rest of the models, which resulted in a baseline flow peak relative fluid velocity of 2 μm/s in the SAS. Even in the case of reduced baseline fluid flow in the SAS, the time-averaged radial Peclet number (Fig. S4a), PVS fluid position (Fig. S4b) and PVS fluid trajectories (Fig. S4c) were essentially unchanged, suggesting that the temporally asymmetric waveform of functional hyperemia can drive directional fluid flow from the PVS to the ECS. The maximum time-averaged radial Peclet number for the model with smaller pressure difference applied across the SAS was 1.93.

### Sleep can enhance solute penetration into the brain

An attractive hypothesis for the purpose of sleep is to remove waste from the brain (Xie *et al.*, 2013; Kress *et al.*, 2014; Jessen *et al.*, 2015), as neurodegeneration is often preceded by sleep disruptions (Mander *et al.*, 2016; MacEdo, Balouch and Tabet, 2017). Enhanced CSF movement has been observed during slow-wave (non-rapid eye movement) sleep in the brain of mice (Xie *et al.*, 2013) and humans (Fultz *et al.*, 2019). Large oscillations of cerebral blood volume (CBV) (Bergel *et al.*, 2018; Turner *et al.*, 2020b) and increased extracellular volume (Xie *et al.*, 2013), which also occur during sleep, have been proposed as the possible mechanisms for driving increased CSF movement during sleep. To determine the relative contributions of the changes in extracellular volume and larger arterial dilations during sleep to enhancing convection, we compared them individually and together to awake-like vasodilations. Using our particle tracking simulations, we examined the PVS fluid movement predicted by increased extracellular volume and large oscillations of CBV, to simulate the sleep state, and compared the resulting fluid movement in the PVS to that during a simulated awake state. The awake state was simulated with the default parameters in Table 1, and a 20% dilation of the artery with an asymmetric waveform, once every 10 seconds. For the simulated sleep state, the increased extracellular volume was simulated by increasing the fluid volume fraction in the domain representing the brain parenchyma from 0.2 to 0.3. The permeability of the ECS in the model was also increased from 2 × 10^−15^m^2^ to 4 × 10^−15^m^2^ to reflect the increased extracellular volume. The large CBV oscillations observed during sleep were simulated by 40% changes in vessel diameter, once every 10 seconds, with the same asymmetric waveform.

Our models suggest that both the increased extracellular volume (and permeability) and the larger CBV oscillations could contribute to larger PVS fluid movement into the ECS. Fig. 5a shows the trajectories of fluid particles for the awake case at the end of 60 seconds and Fig. 5b shows the distribution of particle positions for 60 seconds. In contrast, the particle trajectories and positions during 60 seconds of simulated sleep (Fig. 5d and 5e respectively) show that sleep enhances PVS fluid exchange with the SAS and the ECS. Sleep increases the amount of fluid entering the ECS from the PVS, and the distance of this fluid penetration into the ECS. To examine the contributions of changes in extracellular volume and amplitude of vasodilation to the PVS fluid movement independently, we plotted the fluid position at the end of 30 seconds of simulations, where only one of these changes were made to the awake case (Fig. 5d). The simulations show that increase in extracellular volume (and ECS permeability) changes PVS fluid entering the ECS, without affecting the fluid exchange between the PVS and the SAS, while the increased amplitude of vasodilation enhanced PVS fluid exchange with both SAS and ECS. If ECS permeability increased further, there was further increase in the directional fluid flow from the PVS to the ECS (Fig. S5).

**Fig. 5:**
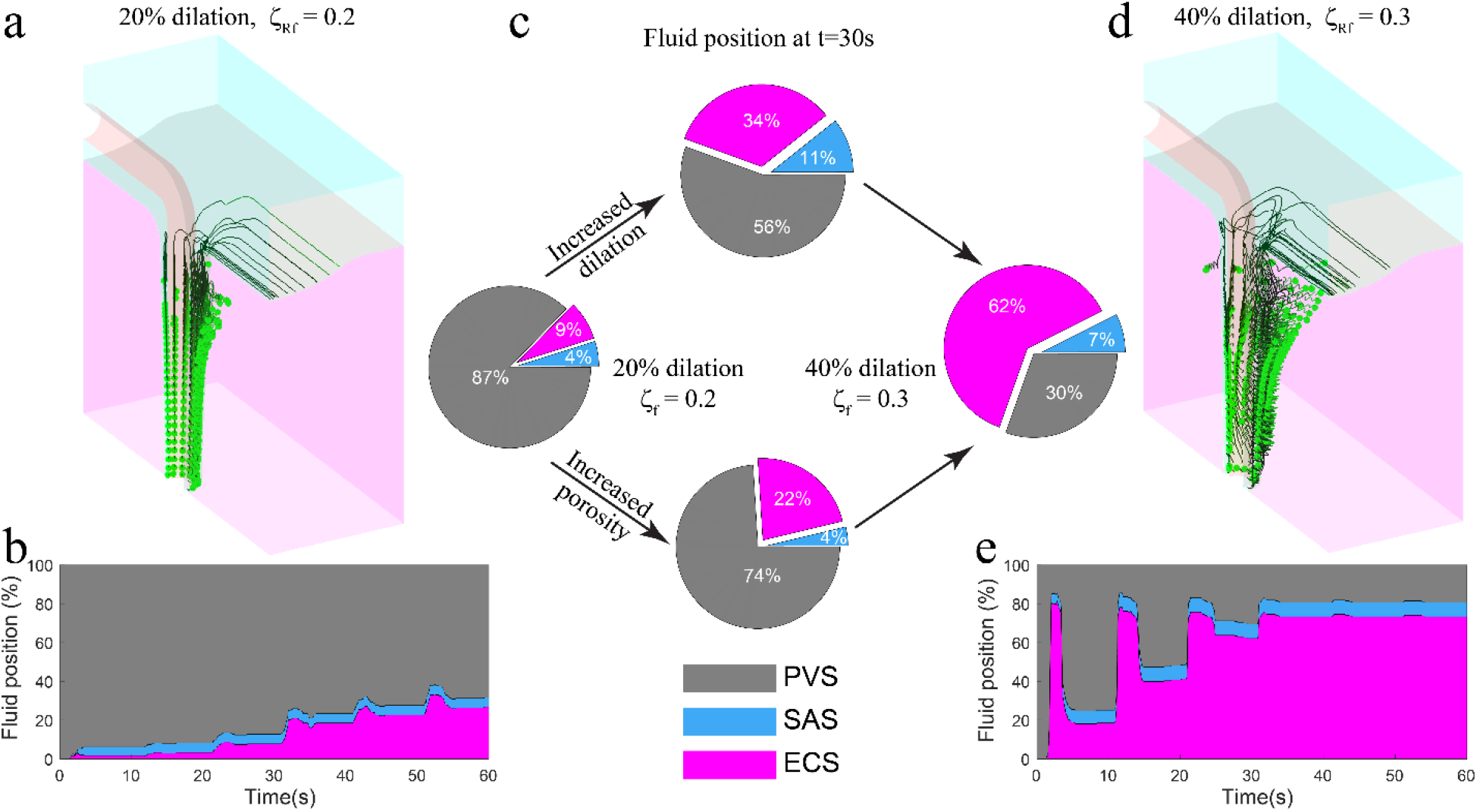
Sleep enhances solute penetration into brain tissue. **a.** The fluid particle trajectories for 60 seconds of simulation for the awake state (20% vessel dilation, once every 10 seconds with 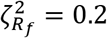). **b.** The fluid particle distribution for the awake state simulation. **c.** The PVS fluid distribution at t=30s for simulated awake and sleep states, along with the cases where only the dilation and porosity are changed. The increase in porosity increased fluid movement into the ECS without affecting the fluid movement into the SAS, while larger dilations increase fluid movement to both SAS and ECS. **d.** The fluid particle trajectories for 60 seconds of simulated sleep. The large amplitude vasodilation during sleep, combined with the increased porosity drives PVS fluid penetration into the brain. **e.** The distribution of fluid position for 60 seconds of simulated sleep. Note: (a) and (b) are repeated from the right side of Fig. 4d and 4c, respectively for better comparison with the sleep state.

Our models also showed that the lower frequency arteriolar diameter changes that occur during sleep (Turner *et al.*, 2020b) could also play a major role in enhancing the directional fluid flow from the PVS to the ECS. We examined the role of frequency of arteriolar dilations in directional fluid flow by changing the dilation time (the time from the start of arteriolar dilation to the return to original size) in the asymmetric dilation waveform (Fig. S6a) and using default values for all other parameters (see Table1). The models showed that amount of PVS fluid exchanged with the ECS increases almost linearly with the increase in dilation time (Fig. S6b). Another way of thinking about the effect of sleep on vasodilation patterns is that the area under the dilation curve (area under curve for arteriolar wall displacement with time) increases during sleep. The effect of the area under the dilation curve on the convective fluid flow through the PVS is demonstrated by our simulations presented in the previous sections (Figs. 3 and 4), which showed that for the same peak vasodilation amplitude, a temporally asymmetric waveform (with nearly two times the area under displacement curve as the symmetric waveform) can drive larger PVS fluid flow into the ECS compared to a symmetric waveform. To further investigate how vasodilation amplitude and area under dilation curve affect PVS fluid flow, we performed simulations with both asymmetric and symmetric dilation waveforms of different peak amplitudes (Fig. S7). The simulations showed that for the same dilation amplitude, the directional fluid flow is appreciably affected by the waveform of the dilation (Fig. S7a,b), while for the same area under dilation curve the directional fluid flow drive by arteriolar dilations is mostly unaffected by the dilation waveform (Fig. S7c,d). Therefore, the larger area under dilation curve observed during sleep could also play an important role in directional fluid flow into the ECS through the PVS.

## Discussion

Here, we simulated fluid flow in a poroelastic model of the brain during the dilations of penetrating arterioles. We found that temporally asymmetric vasodilation drove directed fluid flow, with fluid flow from the PVS into the ECS during dilation, and fluid flows from SAS into the PVS during the return to baseline diameter. This could explain the importance of the speedup of vasodilation by noradrenergic stimulation (Bekar, Wei and Nedergaard, 2012), and how its slowing with age (Handwerker *et al.*, 2007) could contribute to lower solute clearance from the brain. Moreover, given that the brain is oversupplied with oxygen, and that the oxygen changes during functional hyperemia exceed the increased oxygen demand due to neural activity (Leithner and Royl, 2014; Zhang *et al.*, 2019), it is possible that driving fluid movement through the PVS is the main physiological purpose of functional hyperemia. Our poroelastic models showed that the shape, size and frequency of vasodilation, along with the permeability of the ECS are important factors that influence the amount of directional fluid flow from the PVS to the ECS. Based on the results of our simulations, the increased solute transport in the brain during sleep could be attributed to the increased ECS volume, along the large-amplitude, low-frequency vasodilation observed during sleep. While the dilation of arterioles associated with “fidgeting” motions (Drew, Winder and Zhang, 2019; Drew *et al.*, 2020) and exercise (Gao, Greene and Drew, 2015; Gao and Drew, 2016) will help clear waste in the awake brain, it will not be as effective as the larger dilations and porosity changes during sleep.

Several studies have suggested that heartbeat-driven pulsations of arteries and arterioles can cause directional fluid flow through the PVS of arterioles (J. J. Iliff *et al.*, 2013; Bedussi *et al.*, 2017; Mestre, Tithof, *et al.*, 2018; Daversin-Catty *et al.*, 2020). It is possible that heartbeat-driven pulsations, which have a temporally asymmetric waveform (Fujikura *et al.*, 2007; Mestre, Tithof, *et al.*, 2018), can cause directional fluid flow through the PVS of penetrating arterioles. However, issues concerning the choice of boundary conditions and the computational cost of the simulations need to be properly addressed to simulate the fluid flow driven by heartbeat pulsations in a poroelastic brain. While displacement boundary conditions are apt for simulating functional hyperemia, which is a large and active motion of smooth muscle cells and can occur even in brain slices with no perfusion pressure in arteries (Filosa *et al.*, 2006; LeMaistre Stobart *et al.*, 2013; Du, Stern and Filosa, 2015), it is unclear if heartbeat pulsations should be simulated by pressure or displacement boundary conditions, because pulsations are a direct result of pressure changes in the arteriolar lumen. The small scale of the heartbeat pulsations (Bedussi *et al.*, 2017; Mestre, Tithof, *et al.*, 2018; Raghunandan *et al.*, 2021) also makes it hard to tease out the effect of pressure and displacement of the arterial wall. The choice of pressure/displacement boundary conditions could result in widely different predictions in our models, which showed that poroelastic mixtures have very distinct flow characteristics when subjected to pressure and displacement boundary conditions (Fig. S5). The boundary conditions at the end of the SAS in the model also need to be reconsidered. The PVS of arterioles in the cerebral cortex is connected to the PVS of larger arteries, which include major branches of the middle cerebral artery (Bedussi *et al.*, 2017; Mestre, Tithof, *et al.*, 2018; Min Rivas *et al.*, 2020) that also pulsate at heartrate. To understand fluid flow in the PVS of penetrating arterioles, the model needs to include coupled fluid chambers representing the fluid flow in the PVS of large arteries, at the SAS. Another important concern for accurately modeling pulsation-driven fluid flow is the computational cost of simulations. To simulate the frequency response of the model subject to heartbeat pulsations, we need to achieve a state where the change in variables from cycle to cycle is minimal. Since all our simulations have an initial condition where all the variables are set to zero, we need to simulate several cycles of pulsation to achieve the frequency response. In our fluid dynamic (Kedarasetti, Drew and Costanzo, 2020) and fluid-structure interaction (Kedarasetti *et al.*, 2020) models, the frequency response was achieved by slowly ramping up the pulsation amplitude and simulating 20 cycles of pulsation, after which the cycle-to-cycle change in variables was less than 0.1%. Simulating 20 heartbeat cycles with our current 3D poroelastic models, which used a direct solver with 1.3 million unknowns, would be prohibitively expensive for our computational architecture.

There are several ways that the model could be improved. The problem with the computational cost of the model could be addressed by implementing a stabilization techniques (Masud and Hughes, 2002; Olshanskii *et al.*, 2009; Masud and Truster, 2013) for the incompressible poroelastic model, which would allow the usage of first-order interpolations for displacements and velocities, thereby reducing the number of unknowns. A more realistic geometry of the PVS could be used in the model to understand how factors like eccentricity of the PVS affect fluid flow (Tithof *et al.*, 2019). Diffusion equations can be added to the model to simulate tracer infusion experiments more faithfully by including the physics of diffusion and altered mobility of solutes in porous fluid spaces (Sykova *et al.*, 2008). Reduced-order models of the geometry simulated in this study could be used to simulate larger regions of the cortex to better understand the factors influencing large variations in CSF flow observed during exercise (von Holstein-Rathlou, Petersen and Nedergaard, 2018) and sleep (Xie *et al.*, 2013; Fultz *et al.*, 2019).

Our results could shed a new light on some aspects of the glymphatic hypothesis of solute transport in the brain. There has been controversy whether diffusion or convection dominates in the brain (Iliff *et al.*, 2012; Nedergaard, 2013). Our models suggest that directional transport of solutes into the brain is possible, but that it requires the active dilation and constriction of arteries to generate the movement. The model might also explain the controversy over glymphatic flow, in terms of the differences in solute transport in the brain depending on the anesthetic state (Gakuba *et al.*, 2018). Depending on anesthetic state, there may or may not be spontaneous arterial dilations, and without dynamic changes in arterial diameter, there will be much less convective fluid flow in the brain. For example, the lower solute transport seen under isouflurane anesthesia as compared to ketamine/xylazine anesthesia (Hablitz *et al.*, 2019) could be explained in part by the fact that isoflurane is a strong vasodilator, and can occlude vasodilation events (Knutsen, Mateo and Kleinfeld, 2016). Our models could also explain the role of Aqp4 in driving solute transport through the PVS. Knockouts of Aqp4 and α-Syntrophin genes could result in lower permeability at the PVS-ECS interface (Furman *et al.*, 2003; Hoddevik *et al.*, 2017), which could contribute the slower solute transport observed in the brains of in Aqp4 and α-Syntrophin knockout mice (Iliff *et al.*, 2012; Mestre, Hablitz, *et al.*, 2018). In contrast to the fluid pathway from the arteriolar PVS to the venular PVS proposed by the glymphatic hypothesis, our model suggests a pathway of fluid circulation, into the ECS through the PVS of arterioles and out through the surface of the brain. Although the model does not simulate the PVS of venules, the high flow resistance of the ECS and the small size of the PVS of venules compared to arteriolar PVS (Vinje, Bakker and Rognes, 2021) suggest that the path of least resistance for fluid flow out of the ECS is through the brain surface, as predicted by our models. This pathway of fluid circulation would explain why dyes injected into the CSF linger in the PVS of venules (Iliff *et al.*, 2012), where the fluid movement would be minimal, as veins do not dilate in anesthetized animals (Hillman *et al.*, 2007; Drew, Shih and Kleinfeld, 2011).

## Methods

### Model Geometry

The geometry was created using Autodesk Inventor 2020 (San Rafael, Ca.). The two domains (see Fig. 2a), one representing the fluid-filled spaces (SAS and PVS) and one representing the brain parenchyma, were created as separate parts. An assembly was created by matching the two parts, and the assembly was exported into a standard exchange format (.step file), so that it can be accessed by any 3D CAD and meshing software. The step files are available on GitHub (https://github.com/kraviteja89/poroelastic3DPVS).

For the portion representing the fluid-filled spaces, the segment representing the PVS was created by using the loft function between two annular sections 150 μm apart along the *z*-axis. The cross section representing the SAS was created at y = −100 μm and extruded to y = 100 μm. The outer surface of the intersection between the two solids was smoothened by using a fillet of radius 7 μm. The cavity representing the arteriole passing through the SAS was created by using the sweep function on a circular face along a path including a straight line and an arc to connect to the surface and penetrating segments of the arteriole. The solid was split at the *yz*-plane (*x* = 0).

For the part representing the brain tissue, a block of size 160 μm × 200 μm × 170 μm was created by extruding a rectangular face. A cut was made by extruding a negative volume based on the bottom face of the SAS. Another cut was made by using the loft function on two circles representing the outer wall of the PVS. A fillet was made at the intersection of the two faces. The solid was split at the *yz*-plane (*x* = 0).

### Meshing

A custom tetrahedral mesh was generated for the geometry using Altair Hypermesh. The mesh is shown in Fig. 2b. A hexahedral mesh was first created for the PVS surrounding the penetrating segment of the arteriole, with 4 elements along the width of the PVS, 16 elements along the half circumference, and an element height of 3 μm near the surface and 6mm at 150 μm below the surface. Two layers of hexahedron elements of width 1.5 and 2.5 μm perpendicular to the surfaces were created at the interface between the two parts of the geometry, the arteriolar wall in the SAS and the top surface of the dura (*z* = 200 μm). The hexahedral elements were created to control the mesh shape at the interface between the two parts of the geometry and the boundaries where no-slip boundary conditions were applied. These hexahedrons were split into tetrahedrons and controlled triangular meshes were created on the remaining surfaces of each part. The triangular faces of the existing tetrahedrons, along with the triangular meshes on the remining surfaces were used to generate a tetrahedral volume meshes. Quality of mesh was maintained by setting a minimum tet collapse (1.24 × ratio of distance between a node from the opposite triangular face to the area of the face) of 0.15. The mesh was exported into the Nastran format (.nas), which was imported into COMSOL Multiphysics.

### Model Formulation

A poroelastic model (Costanzo and Miller, 2017) was used to simulate fluid flow through the SAS, PVS and the ECS, along with the deformations of the connective and parenchymal tissue. The model is divided into two domains, one representing the fluid-filled PVS and SAS (Ω^1^) and the other representing the parenchymal tissue (Ω^2^), as described in the model geometry section. In each domain, we solve for five unknowns 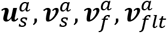 and *p*^*a*^ (4 vectors and 1 scalar) representing the solid displacement, solid velocity, fluid velocity, filtration velocity and pore pressure respectively. The superscript, *a* = 1, 2 represents the domain. In both domains, an arbitrary Lagrangian–Eulerian (ALE) finite element formulation of poroelasticity, based on mixture theory was implemented to simulate an incompressible hyperelastic skeleton saturated with an incompressible fluid. The development of the formulation is explained in detail by Costanzo and Miller (Costanzo and Miller, 2017). The key equations of the formulation are described below.

#### Kinematics

The ALE formulation is written in the coordinates of the undeformed solid skeleton 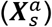. The domains in the reference (undeformed solid) frame are represented by 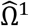 and 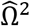. If the deformed configuration is given by 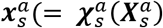, where 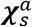 is a smooth map describing the transformation from the deformed state to the reference configuration), the displacement 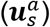, deformation gradient 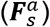 and Jacobian determinant 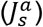 of the motion are given by Eq. (1). In Eq. (1), ∇ is the gradient operator with respect to the reference coordinates, ***I*** is the identity tensor, and det is the determinant.

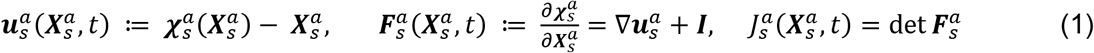

Since the equations are written in the material particle coordinates of the solid skeleton, the relation between solid displacement and solid velocity is given by Eq. (2).

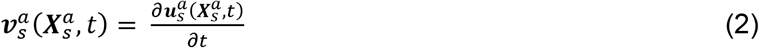

At each point, the volume fraction of the solid and fluid are given by 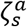 and 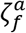 respectively and the mass densities are given by 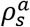 and 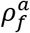 respectively. The mass densities are related to the true densities (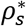 and 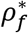) of the solid and fluid phases by Eq. (3). The true mass density of a component of the mixture is defined as the mass density of that component in its single-phase state. Note that the true densities are constants for incompressible phases and hence the superscript is omitted. Since the solid is fully saturated by the fluid, the sum of their volume fractions is unity, which is the constraint shown in Eq. (4).

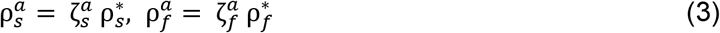

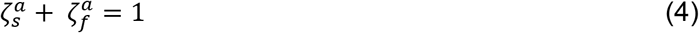

As the solid deforms, the volume fractions of the phases evolve continuously. The relation between, 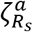, the volume fraction of the solid in the undeformed reference configuration and the actual volume fractions of the phases are given by Eq. (5).

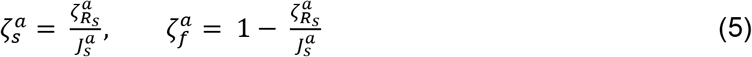

The filtration velocity 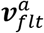 is defined as the velocity of the fluid relative to the solid skeleton scaled by the volume fraction of the fluid, as shown in Eq. (6).

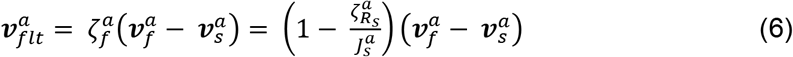

In the absence of chemical reactions, the incompressibility constraint is given by the zero divergence of the volume averaged velocity, thus yielding the constraint shown in Eq. (7), where div and grad are the divergence and gradients with respect to the deformed coordinates 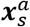, while the colon : denotes the inner product of tensors. Using the chain rule and Eq. (6), the incompressibility constraint, written in terms of quantities defined in the ALE coordinates, takes the form shown in Eq. (8), where ***A***^−1^ and ***A***^*T*^ are the inverse and transpose operations respectively on the tensor ***A***:

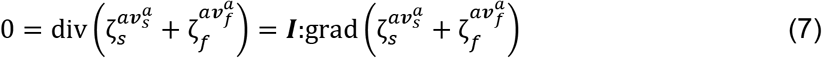

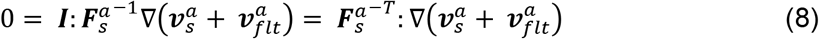

#### Constitutive assumptions and Momentum balance

The solid skeleton is modeled as an isotropic incompressible Neo-Hookean solid and the fluid flow is modeled by incompressible Darcy-Brinkman law for flow through porous solids. The total Cauchy stress of the mixture is given by Eq. (9), where, 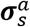 is the elastic contribution due to the strain energy density (Ψ_*s*_), while 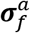 accounts for the Brinkman dissipation. In Eq. (10), the shear modulus of the solid skeleton for domain *a* is given by 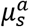. The strain energy density in Eq. (10) appears similar to that of a compressible solid, which is a valid choice of constitutive model because even though the pure solid constituent is incompressible, the solid skeleton of the porous solid can be compressed (Treloar, 1975; Rajagopal, Wineman and Gandhi, 1986) and it has the convenience of yielding the expression 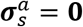 in the reference configuration. In Eq. (11) *μ_f_* is the dynamic viscosity of the fluid. It is important to note that the stresses in Eqs. (9)–(11) are defined in terms of the deformed coordinates.

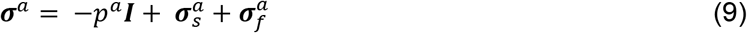

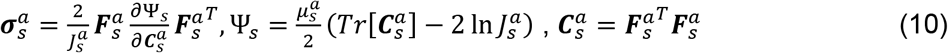

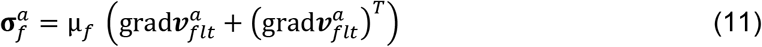

The momentum equations can be written in the ALE coordinates based on the stresses defined in Eqs. (9)–(11) and the chain rule for transforming the spatial gradients. Equations (12) and (13) are the momentum equations for the solid and fluid components of the mixture respectively. The definitions of 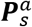 and 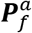 are given by Eqs. (14) and (15), respectively.

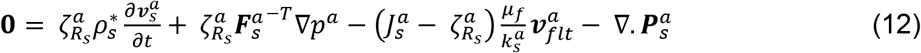

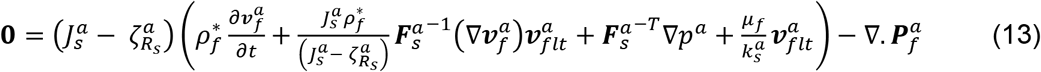

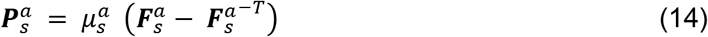

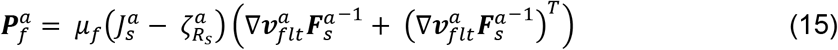

Within each domain, we have five equations, Eqs. (2), (6), (8), (12), and (13), that govern the spatiotemporal evolution of the five primary unknowns.

#### Interface Conditions

At the interface of the mostly fluid-filled spaces *i.e.*, the SAS and the PVS with the brain parenchyma, there is a sharp change (mathematically, a possible jump discontinuity) in the volume fraction of the fluid (porosity) and the composition of the solid skeleton. Therefore, there is a sharp change in the fluid permeability and elastic modulus which could result in a sharp change in the fluid velocity and the distribution of traction between the solid and fluid components at the interface. To deal with the sharp change, the mass and traction continuity at the interface were implemented through special boundary conditions called jump conditions (dell’Isola, Madeo and Seppecher, 2009; Shim and Ateshian, 2021; Shim *et al.*, 2021).

The solid phases in both the domains are in contact with each other at the interface and therefore the solid displacement and the velocity are continuous across the interface, as indicated in Eq. (17). For an incompressible fluid (fluid with constant true density), the mass conservation for the fluid across the interface dictates that the component of the filtration velocity normal to the interface should be continuous across the boundary. By considering the limiting case as 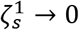, Hou et al. (Hou *et al.*, 1989) showed that the no-slip condition for both extremes at the other end, 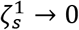 and 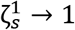) are valid when the tangential component of filtration velocity is continuous across the interface. The continuity of both tangential and normal components of the filtration velocities implies the continuity of filtration velocity across the interface, indicated in Eq. (17):

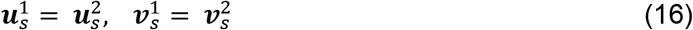

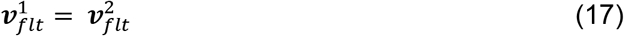

Equation (18) states the condition that the *total* traction force across the interface be continuous. In Eq. (18), ***n***^1^ and ***n***^2^ are the unit outward normal to the domains Ω^1^ and Ω^2^ respectively. Additionally, we assume that the ratio of tractions on each phase is equal to the ratio of the volume fractions of the phase. This assumption is stated in Eqs. (19) and (20).

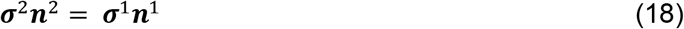

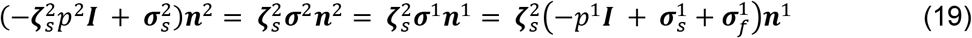

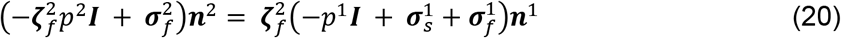

Equations (18)–(20) are written in the deformed configuration. The unit outward normal to Ω^*a*^, ***n***^*a*^, is related to the unit outward normal to 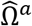 (the undeformed domain), 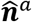, according to the relation in Eq. (21). Using Eqs. (14), (15), and (21), the traction conditions at the interface can then be rewritten in ALE coordinates as shown in Eqs. (22) and (23).

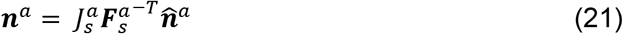

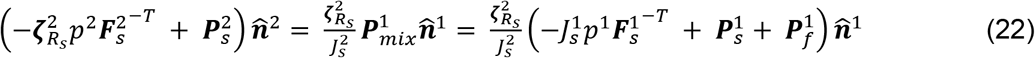

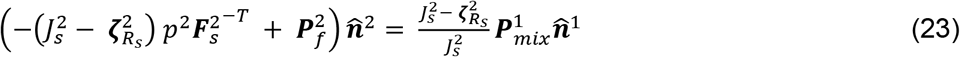

#### Boundary conditions

For the segment of the arteriole in the SAS (z > 150 μm), the displacement is prescribed against the direction of the outward normal (Eq. (24)). For the segment of the arteriole (z ≤ 150 μm), the displacement is prescribed along the radially outward direction as shown in Eq. (25) where *R*_0_ is the nominal radius of the vessel (see Table 1). The solid velocity on the arteriolar wall is the partial time derivative of the prescribed displacement, shown in Eq. (26), where 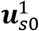 is the displacement prescribed by Eqs. (24) and (25). The no-slip condition is implemented by setting the filtration velocity to zero.

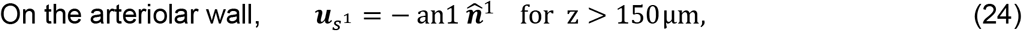

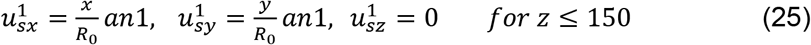

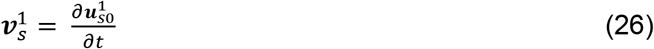

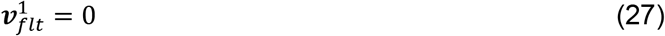

At the bottom face of the fluid-filled domain 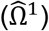, the solid displacement and velocity in the *z* direction were set to zero (Eq. (28)). On the fluid phase, a flow-dependent traction (flow resistance) boundary condition was used. The flow resistance at the bottom end of the PVS was set to 10 times the resistance of an annular region with the permeability of the PVS, inner radius of 7.5 μm (*R*_2_) and a width of 5.5 μm (*W*_2_). In Eq. (29), *L*_*a*_ is the height of the PVS segment (150 μm) and *Q*_1_ is the flowrate through the bottom face calculated by the integral over the bottom face 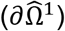 defined in Eq. (30).

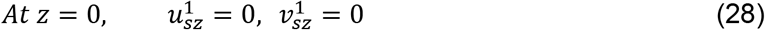

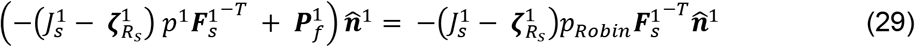

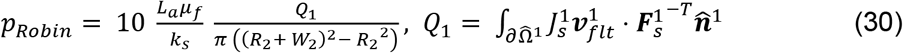

The circulation of CSF in the SAS was simulated by applying a small pressure difference across the ends of the SAS on the fluid component (green and blue faces in Fig. 2a). The solid displacement and velocity in the y directions were set to zero at the ends of the SAS.

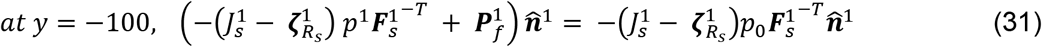

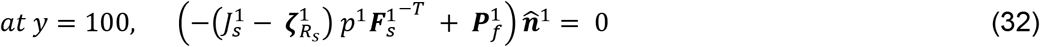

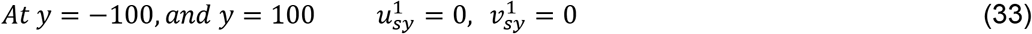

At the top surface of the SAS, which represents the dura, all the velocities and displacements were set to zero.

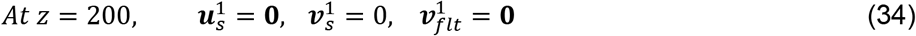

At *x* = 0 and *x* = 80, symmetry boundary conditions were used, where the solid displacement, velocity and filtration velocity normal to the surface were set to zero.

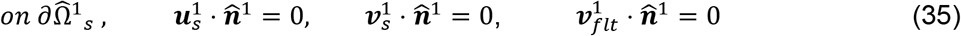

At all the surfaces on the domain representing the brain tissue 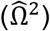, other than the interface with the fluid-filled domain, the normal components of displacements and velocities were set to zero. At the plane of symmetry (*x* = 0), the boundary condition is self-explanatory. The use of the boundary condition at the side-facing surfaces (*y* = −100, *y* = 100 and *x* = 80), the boundary condition represents the assumption that the region being modeled is surrounded by similar blocks of the brain tissue experiencing vasodilation. On the bottom surface, a flow resistance boundary condition, similar to the one presented in Eq. 29 was not applied, as it would set a constant traction force on the whole surface.

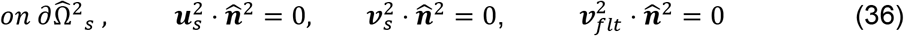

#### Finite element implementation

The overall initial-boundary value problem formulated in the previous section is very complex and nonlinear. To the authors’ knowledge, there are no formal proofs available in the literature to guide us in the formulation of a well-posed weak problem with minimum smoothness requirements for the various unknown fields under reasonable smoothness assumptions on the problem’s data. Consequently, here we make assumptions as to the functional spaces that we wish to have available to make the writing of the problem meaningful. Our numerical results indicate that our finite element implementation of the problem is adequate (Costanzo and Miller, 2017). However, we wish to be clear that we are in no position to offer rigorous proofs as yet on the well-posedness of the problem. First, we consider vector functional spaces 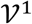 and 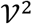, along with scalar functional spaces 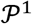 and , 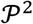 that are defined as follows.

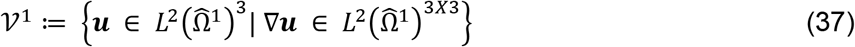

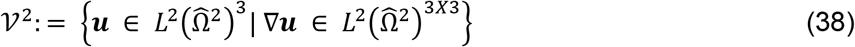

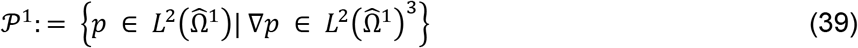

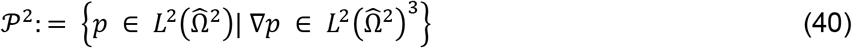

Next, we define the solution spaces for 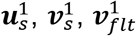 as subsets of 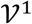. The boundaries of the domain 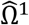 are divided into four subsets such that, 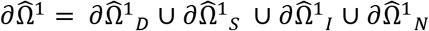, where 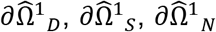 are the surfaces where Dirichlet (vessel wall and skull), symmetry (*x* = 0 and *x* = 80) and Neumann (*y* = −100, *y* = 100 and *z* = 0) boundary conditions are prescribed on the fluid phase, while 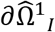 is the interface boundary between the two domains.

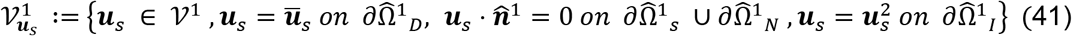

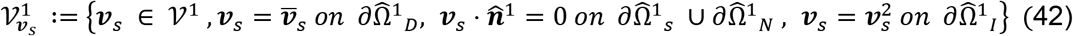

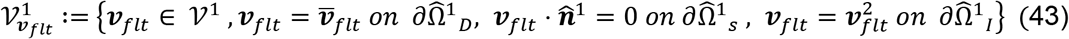

We also define the companion spaces for the test functions as follows.

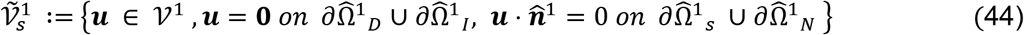

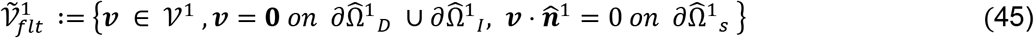

The solutions and test functions for 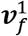 and *p*^1^ are taken from 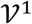 and 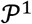 respectively.

For 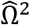, the boundary is divided into two non-intersecting subsets, 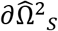 and 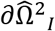, representing the boundaries with the symmetry and interface conditions respectively. The solution and test functions for 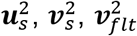 belong to the same functional space (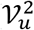 in Eq. 46). The solutions and test functions for 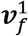 and *p*^2^ are taken from 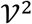 and 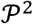 respectively.

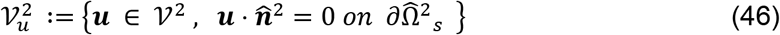

To simplify the weak form, we define the following notation, where *a* = 1, 2, represents the domain and 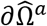 represents the boundary.

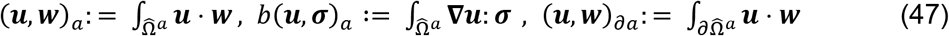

The weak form of the problem can be written as follows:

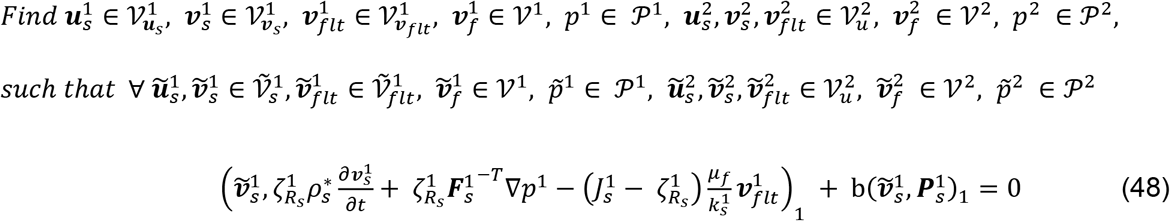

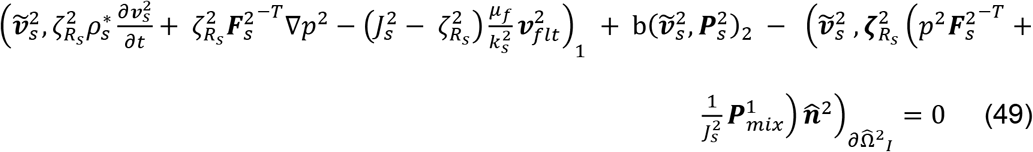

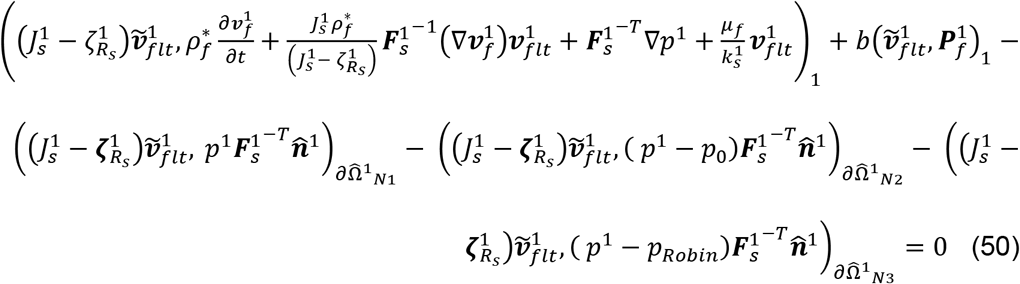

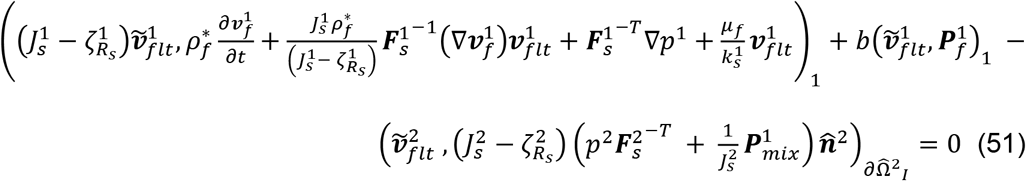

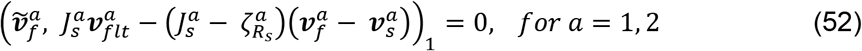

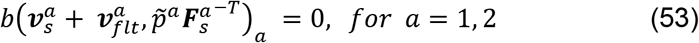

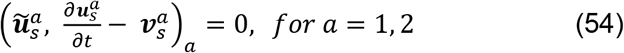

In Eq. 50, 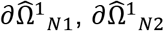 and 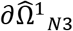 are the boundaries to 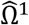 at *y* = −100, *y* = 100 and *z* = 0, respectively.

These weak form equations were converted to their component form using Wolfram Mathematica. The equations were implemented using the weak form PDE module in COMSOL Multiphysics. A mixed-finite element model was used with second order Lagrange polynomials for all variables except pressure, which used a first order Lagrange polynomial. The initial conditions were set to zero value for all variables. A baseline time-dependent problem was solved, where the magnitude of the traction on 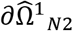 was ramped from 0 to *p*_0_ in 0.1 seconds, with no arteriolar dilation. The baseline problem was run for 0.5 seconds, to reach a steady state (the changes in filtration velocity less than 0.01%). The simulations with vasodilation were then performed with the initial conditions set to the last timestep of the baseline model, and the outputs were saved for 201 time points between 0.5 and 10.5. Second order backward difference formula (BDF), with a timestep of 0.0025s was used to solve the time-dependent problems.

### Fluid particle tracking

For tracking the motion of fluid, we need the fluid particle velocities in the computational frame. Therefore, we used the fluid velocities and the displacement fields calculated using the finite element model and calculated the fluid particle velocity in the computational frame. The equation for fluid particle velocity in the computational frame (Eq. 55) was derived in our previous publication (Kedarasetti, Drew and Costanzo, 2020) and is valid for both domains. The fluid velocities for both the domains were exported into a text format using COMSOL Multiphysics for all the points on the computational grid for the 201 time points to be used for fluid particle tracking.

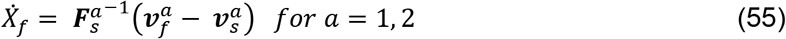

A grid of equally spaced points (Fig. 3a) was created using Altair Hypermesh. Similarly, grids and meshed were created for the boundaries of the two domains and the surface representing the arteriolar wall. The grids and meshes were converted to text format using Microsoft Excel.

The fluid particle velocities, along with the grids and meshes were imported into MATLAB. The data was repeated along the time axis to calculate the fluid particle trajectories for 60 seconds. The particle trajectories were calculated by interpolating the imported velocity and using a backward Euler integration scheme.

### Peclet numbers

Two Peclet numbers were defined to compare the convective and diffusive transport of solutes driven by vasodilation. The axial Peclet number (*Pe_a_*), was defined based on the flow through the bottom of the PVS (*z* = 0), and represents pumping by arteriolar wall movements similar to peristaltic pumping. In Eq. 56, *Q*_*a*_ and *A*_*a*_ are the flowrate and flow surface area of the bottom surface of the PVS and *D*_*aβ*_ is the diffusion coefficient of amyloid-β.

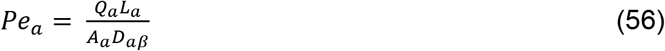

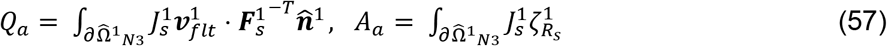

The radial Peclet number was defined to represent solute penetration into the brain parenchyma and was based on the relative velocity of the fluid with respect to the solid in the radial direction into the ECS, in the immediate vicinity of the PVS-ECS interface (Eq. 58). In Eq. 58, 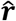 is the unit normal vector in the radially outward direction and *λ* is the tortuosity of the ECS.

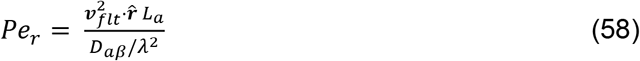

## Acknowledgments

The authors thank Maiken Nedergaard and Douglas Kelley for providing valuable feedback on the manuscript.

## References

Abbott, N. J. et al. (2018) ‘The role of brain barriers in fluid movement in the CNS: is there a “glymphatic” system?’, Acta Neuropathologica, 135(3), pp. 1–21. doi: 10.1007/s00401-018-1812-4.

Adams, M. D. et al. (2018) ‘The pial vasculature of the mouse develops according to a sensory-independent program’, Scientific Reports, 8(1), pp. 1–12. doi: 10.1038/s41598-018-27910-3.

Asgari, M., De Zélicourt, D. and Kurtcuoglu, V. (2015) ‘How astrocyte networks may contribute to cerebral metabolite clearance’, Scientific Reports, 5, pp. 1–13. doi: 10.1038/srep15024.

Asgari, M., De Zélicourt, D. and Kurtcuoglu, V. (2016) ‘Glymphatic solute transport does not require bulk flow’, Scientific Reports, 6, pp. 1–11. doi: 10.1038/srep38635.

Barber, T. W., Brockway, J. A. and Higgins, L. S. (1970) ‘The density of tissues in and about the head’, Acta neurologica scandinavica, 46(1), pp. 85–92.

Bedussi, B. et al. (2017) ‘Paravascular spaces at the brain surface: Low resistance pathways for cerebrospinal fluid flow’, Journal of Cerebral Blood Flow & Metabolism, p. 0271678X1773798. doi: 10.1177/0271678X17737984.

Bekar, L. K., Wei, H. S. and Nedergaard, M. (2012) ‘The locus coeruleus-norepinephrine network optimizes coupling of cerebral blood volume with oxygen demand’, Journal of Cerebral Blood Flow \& Metabolism, 32(12), pp. 2135–2145.

Bergel, A. et al. (2018) ‘Local hippocampal fast gamma rhythms precede brain-wide hyperemic patterns during spontaneous rodent REM sleep’, Nature Communications, 9(1). doi: 10.1038/s41467-018-07752-3.

Bilston, L. E. et al. (2003) ‘Arterial pulsation-driven cerebrospinal fluid flow in the perivascular space: a computational model’, Computer Methods in Biomechanics & Biomedical Engineering, 6(4), pp. 235–241.

Blinder, P. et al. (2013) ‘The cortical angiome: An interconnected vascular network with noncolumnar patterns of blood flow’, Nature Neuroscience, 16(7), pp. 889–897. doi: 10.1038/nn.3426.

Bowen, R. M. (1976) ‘Theory of mixtures in Continuum physics III, Ed. AC Eringen’. Academic Press, New York.

Bowen, R. M. (1980) ‘Incompressible porous media models by use of the theory of mixtures’, International Journal of Engineering Science, 18(9), pp. 1129–1148. doi: 10.1016/0020-7225(80)90114-7.

Bradbury, M. W. B., Cserr, H. F. and Westrop, R. J. (1981) ‘Drainage of cerebral interstitial fluid into deep cervical lymph of the rabbit’, American Journal of Physiology - Renal Fluid and Electrolyte Physiology, 9(4), pp. 329–336. doi: 10.1152/ajprenal.1981.240.4.f329.

Budday, S. et al. (2017) ‘Mechanical characterization of human brain tissue’, Acta Biomaterialia, 48, pp. 319–340. doi: 10.1016/j.actbio.2016.10.036.

Budday, S. et al. (2019) Fifty Shades of Brain: A Review on the Mechanical Testing and Modeling of Brain Tissue, Archives of Computational Methods in Engineering. Springer Netherlands. doi: 10.1007/s11831-019-09352-w.

Coles, J. A., Stewart-Hutchinson, P. J., et al. (2017) ‘The mouse cortical meninges are the site of immune responses to many different pathogens, and are accessible to intravital imaging’, Methods, 127, pp. 53–61. doi: 10.1016/j.ymeth.2017.03.020.

Coles, J. A., Myburgh, E., et al. (2017) ‘Where are we? The anatomy of the murine cortical meninges revisited for intravital imaging, immunology, and clearance of waste from the brain’, Progress in Neurobiology, 156, pp. 107–148. doi: 10.1016/j.pneurobio.2017.05.002.

Costanzo, F. and Miller, S. T. (2017) ‘An arbitrary Lagrangian–Eulerian finite element formulation for a poroelasticity problem stemming from mixture theory’, Computer Methods in Applied Mechanics and Engineering, 323, pp. 64–97. doi: 10.1016/j.cma.2017.05.006.

Cserr, H. F., Harling-Berg, C. J. and Knopf, P. M. (1992) ‘Drainage of Brain Extracellular Fluid into Blood and Deep Cervical Lymph and its Immunological Significance’, Brain Pathology, 2(4), pp. 269–276. doi: 10.1111/j.1750-3639.1992.tb00703.x.

Das, A., Murphy, K. and Drew, P. J. (2021) ‘Rude mechanicals in brain haemodynamics: non-neural actors that influence blood flow’, Philosophical Transactions of the Royal Society B, 376(1815), p. 20190635.

Daversin-Catty, C. et al. (2020) ‘The mechanisms behind perivascular fluid flow’, PLoS ONE, 15(12 December), pp. 1–20. doi: 10.1371/journal.pone.0244442.

dell’Isola, F., Madeo, A. and Seppecher, P. (2009) ‘Boundary conditions at fluid-permeable interfaces in porous media: A variational approach’, International Journal of Solids and Structures, 46(17), pp. 3150–3164. doi: 10.1016/j.ijsolstr.2009.04.008.

Drew, P. J. et al. (2010) ‘Chronic optical access through a polished and reinforced thinned skull.’, Nature methods, 7(12), pp. 981–984. doi: 10.1038/nmeth.1530.

Drew, P. J. (2019) ‘Vascular and neural basis of the BOLD signal’, Current Opinion in Neurobiology, 58, pp. 61–69. doi: 10.1016/j.conb.2019.06.004.

Drew, P. J. et al. (2020) ‘Ultra-slow oscillations in fMRI and Resting-State connectivity: neuronal and vascular contributions and technical confounds’, Neuron.

Drew, P. J., Shih, A. Y. and Kleinfeld, D. (2011) ‘Fluctuating and sensory-induced vasodynamics in rodent cortex extend arteriole capacity’, Proceedings of the National Academy of Sciences, 108(20), pp. 8473–8478. doi: 10.1073/pnas.1100428108.

Drew, P. J., Winder, A. T. and Zhang, Q. (2019) ‘Twitches, Blinks, and Fidgets: Important Generators of Ongoing Neural Activity’, Neuroscientist, 25(4), pp. 298–313. doi: 10.1177/1073858418805427.

Du, W., Stern, J. E. and Filosa, J. A. (2015) ‘Neuronal-Derived Nitric Oxide and Somatodendritically Released Vasopressin Regulate Neurovascular Coupling in the Rat Hypothalamic Supraoptic Nucleus’, Journal of Neuroscience, 35(13), pp. 5330–5341. doi: 10.1523/JNEUROSCI.3674-14.2015.

Filosa, J. A. et al. (2006) ‘Local potassium signaling couples neuronal activity to vasodilation in the brain’, Nature Neuroscience, 9(11), pp. 1397–1403. doi: 10.1038/nn1779.

Fujikura, K. et al. (2007) ‘A novel noninvasive technique for pulse-wave imaging and characterization of clinically-significant vascular mechanical properties in vivo’, Ultrasonic Imaging, 29(3), pp. 137–154. doi: 10.1177/016173460702900301.

Fultz, N. E. et al. (2019) ‘Coupled electrophysiological, hemodynamic, and cerebrospinal fluid oscillations in human sleep’, Science, 366(6465), pp. 628–631. doi: 10.1126/science.aax5440.

Furman, C. S. et al. (2003) ‘Aquaporin-4 square array assembly: Opposing actions of M1 and M23 isoforms’, Proceedings of the National Academy of Sciences of the United States of America, 100(23), pp. 13609–13614. doi: 10.1073/pnas.2235843100.

Gagnon, L. et al. (2015) ‘Quantifying the microvascular origin of bold-fMRI from first principles with two-photon microscopy and an oxygen-sensitive nanoprobe’, Journal of Neuroscience, 35(8), pp. 3663–3675. doi: 10.1523/JNEUROSCI.3555-14.2015.

Gakuba, C. et al. (2018) ‘General anesthesia inhibits the activity of the “glymphatic system”’, Theranostics, 8(3), pp. 710–722. doi: 10.7150/thno.19154.

Gao, X. Y. and Drew, X. P. J. (2016) ‘Effects of Voluntary Locomotion and Calcitonin Gene-Related Peptide on the Dynamics of Single Dural Vessels in Awake Mice’, 36(8), pp. 2503–2516. doi: 10.1523/JNEUROSCI.3665-15.2016.

Gao, Y. R., Greene, S. E. and Drew, P. J. (2015) ‘Mechanical restriction of intracortical vessel dilation by brain tissue sculpts the hemodynamic response’, NeuroImage, 115, pp. 162–176. doi: 10.1016/j.neuroimage.2015.04.054.

Hablitz, L. M. et al. (2019) ‘Increased glymphatic influx is correlated with high EEG delta power and low heart rate in mice under anesthesia’, Science Advances, 5(2). doi: 10.1126/sciadv.aav5447.

Handwerker, D. A. et al. (2007) ‘Reducing vascular variability of fMRI data across aging populations using a breathholding task’, Human brain mapping, 28(9), pp. 846–859.

Hardy, J. A. and Higgins, G. A. (1992) ‘Alzheimer’s disease: the amyloid cascade hypothesis’, Science, 256(5054), pp. 184–186.

He, Y. et al. (2018) ‘Ultra-slow single-vessel BOLD and CBV-based fMRI spatiotemporal dynamics and their correlation with neuronal intracellular calcium signals’, Neuron, 97(4), pp. 925–939.

Hillman, E. M. C. et al. (2007) ‘Depth-resolved optical imaging and microscopy of vascular compartment dynamics during somatosensory stimulation’, Neuroimage, 35(1), pp. 89–104.

Hoddevik, E. H. et al. (2017) ‘Factors determining the density of AQP4 water channel molecules at the brain–blood interface’, Brain Structure and Function, 222(4), pp. 1753–1766. doi: 10.1007/s00429-016-1305-y.

von Holstein-Rathlou, S., Petersen, N. C. and Nedergaard, M. (2018) ‘Voluntary running enhances glymphatic influx in awake behaving, young mice’, Neuroscience Letters, 662(October 2017), pp. 253–258. doi: 10.1016/j.neulet.2017.10.035.

Holter, K. E. et al. (2017) ‘Interstitial solute transport in 3D reconstructed neuropil occurs by diffusion rather than bulk flow’, Proceedings of the National Academy of Sciences, p. 201706942. doi: 10.1073/pnas.1706942114.

Horton, N. G. et al. (2013) ‘In vivo three-photon microscopy of subcortical structures within an intact mouse brain’, Nature Photonics, 7(3), pp. 205–209. doi: 10.1038/nphoton.2012.336.

Hou, J. S. et al. (1989) ‘Boundary conditions at the cartilage-synovial fluid interface for joint lubrication and theoretical verifications’, Journal of Biomechanical Engineering, 111(1), pp. 78–87. doi: 10.1115/1.3168343.

Iadecola, C. (2017) ‘The Neurovascular Unit Coming of Age: A Journey through Neurovascular Coupling in Health and Disease’, Neuron, 96(1), pp. 17–42. doi: 10.1016/j.neuron.2017.07.030.

Iliff, J. J. et al. (2012) ‘A Paravascular Pathway Facilitates CSF Flow Through the Brain Parenchyma and the Clearance of Interstitial Solutes, Including Amyloid’, Science Translational Medicine, 4(147), pp. 147ra111–147ra111. doi: 10.1126/scitranslmed.3003748.

Iliff, Jeffrey J. et al. (2013) ‘Brain-wide pathway for waste clearance captured by contrast-enhanced MRI’, Journal of Clinical Investigation, 123(3), pp. 1299–1309. doi: 10.1172/JCI67677.

Iliff, J. J. et al. (2013) ‘Cerebral Arterial Pulsation Drives Paravascular CSF-Interstitial Fluid Exchange in the Murine Brain’, Journal of Neuroscience, 33(46), pp. 18190–18199. doi: 10.1523/JNEUROSCI.1592-13.2013.

Iliff, J. and Simon, M. (2019) ‘CrossTalk proposal: The glymphatic system supports convective exchange of cerebrospinal fluid and brain interstitial fluid that is mediated by perivascular aquaporin-4’, The Journal of Physiology, 0, pp. 1–3. doi: 10.1113/jp277635.

Jessen, N. A. et al. (2015) ‘The Glymphatic System: A Beginner’s Guide’, Neurochemical Research, 40(12), pp. 2583–2599. doi: 10.1007/s11064-015-1581-6.

Jin, B. J. et al. (2013) ‘Aquaporin-4-dependent K+ and water transport modeled in brain extracellular space following neuroexcitation’, Journal of General Physiology, 141(1), pp. 119–132. doi: 10.1085/jgp.201210883.

Kedarasetti, R. T. et al. (2020) ‘Functional hyperemia drives fluid exchange in the paravascular space’, Fluids and Barriers of the CNS, 17(1), pp. 1–25.

Kedarasetti, R. T., Drew, P. J. and Costanzo, F. (2020) ‘Arterial pulsations drive oscillatory flow of CSF but not directional pumping.’, Scientific reports, 10(1), p. 10102. doi: 10.1038/s41598-020-66887-w.

Knutsen, P. M., Mateo, C. and Kleinfeld, D. (2016) ‘Precision mapping of the vibrissa representation within murine primary somatosensory cortex’, Philosophical Transactions of the Royal Society B: Biological Sciences, 371(1705). doi: 10.1098/rstb.2015.0351.

Korogod, N., Petersen, C. C. H. and Knott, G. W. (2015) ‘Ultrastructural analysis of adult mouse neocortex comparing aldehyde perfusion with cryo fixation’, eLife, 4(AUGUST2015), pp. 1–17. doi: 10.7554/eLife.05793.

Kress, B. T. et al. (2014) ‘Impairment of paravascular clearance pathways in the aging brain’, Annals of Neurology, 76(6), pp. 845–861. doi: 10.1002/ana.24271.

Leithner, C. and Royl, G. (2014) ‘The oxygen paradox of neurovascular coupling’, Journal of Cerebral Blood Flow and Metabolism, 34(1), pp. 19–29. doi: 10.1038/jcbfm.2013.181.

LeMaistre Stobart, J. L. et al. (2013) ‘Astrocyte-induced cortical vasodilation is mediated by D-serine and endothelial nitric oxide synthase’, Proceedings of the National Academy of Sciences of the United States of America, 110(8), pp. 3149–3154. doi: 10.1073/pnas.1215929110.

Louveau, A. et al. (2015) ‘Structural and functional features of central nervous system lymphatic vessels’, Nature, 523(7560), pp. 337–341. doi: 10.1038/nature14432.

Ma, Q. et al. (2017) ‘Outflow of cerebrospinal fluid is predominantly through lymphatic vessels and is reduced in aged mice’, Nature Communications, 8(1). doi: 10.1038/s41467-017-01484-6.

MacEdo, A. C., Balouch, S. and Tabet, N. (2017) ‘Is Sleep Disruption a Risk Factor for Alzheimer’s Disease?’, Journal of Alzheimer’s Disease, 58(4), pp. 993–1002. doi: 10.3233/JAD-161287.

Mander, B. A. et al. (2016) ‘Sleep: A Novel Mechanistic Pathway, Biomarker, and Treatment Target in the Pathology of Alzheimer’s Disease?’, Trends in Neurosciences, 39(8), pp. 552–566. doi: 10.1016/j.tins.2016.05.002.

Martinac, A. D. and Bilston, L. E. (2019) ‘Computational modelling of fluid and solute transport in the brain’, Biomechanics and Modeling in Mechanobiology, (0123456789). doi: 10.1007/s10237-019-01253-y.

Massi, F. et al. (2001) ‘Simulation study of the structure and dynamics of the Alzheimer’s amyloid peptide congener in solution’, Biophysical Journal, 80(1), pp. 31–44. doi: 10.1016/S0006-3495(01)75993-0.

Masud, A. and Hughes, T. J. R. (2002) ‘A stabilized mixed finite element method for Darcy flow’, Computer methods in applied mechanics and engineering, 191(39-40), pp. 4341–4370. doi: 10.1016/S0045-7825(02)00371-7.

Masud, A. and Truster, T. J. (2013) ‘A framework for residual-based stabilization of incompressible finite elasticity: Stabilized formulations and F methods for linear triangles and tetrahedra’, Computer Methods in Applied Mechanics and Engineering, 267, pp. 359–399. doi: 10.1016/j.cma.2013.08.010.

Mestre, H., Hablitz, L. M., et al. (2018) ‘Aquaporin-4-dependent glymphatic solute transport in the rodent brain’, eLife, 7, pp. 1–31. doi: 10.7554/elife.40070.

Mestre, H., Tithof, J., et al. (2018) ‘Flow of cerebrospinal fluid is driven by arterial pulsations and is reduced in hypertension’, Nature communications, 9(1), p. 4878. doi: 10.1038/s41467-018-07318-3.

Mestre, H., Mori, Y. and Nedergaard, M. (2020) ‘The Brain’s Glymphatic System: Current Controversies’, Trends in Neurosciences, 43(7), pp. 458–466. doi: 10.1016/j.tins.2020.04.003.

Mihai, L. A. et al. (2017) ‘A family of hyperelastic models for human brain tissue’, Journal of the Mechanics and Physics of Solids, 106, pp. 60–79. doi: 10.1016/j.jmps.2017.05.015.

Min Rivas, F. et al. (2020) ‘Surface periarterial spaces of the mouse brain are open, not porous: Surface periarterial spaces of the mouse brain are open, not porous’, Journal of the Royal Society Interface, 17(172). doi: 10.1098/rsif.2020.0593rsif20200593.

Nedergaard, M. (2013) ‘Neuroscience. Garbage truck of the brain.’, Science (New York, N.Y.), 340(6140), pp. 1529–30. doi: 10.1126/science.1240514.

Neeves, K. B. et al. (2006) ‘Fabrication and characterization of microfluidic probes for convection enhanced drug delivery’, Journal of Controlled Release, 111(3), pp. 252–262. doi: 10.1016/j.jconrel.2005.11.018.

Nishimura, N. et al. (2007) ‘Penetrating arterioles are a bottleneck in the perfusion of neocortex’, Proceedings of the National Academy of Sciences of the United States of America, 104(1), pp. 365–370. doi: 10.1073/pnas.0609551104.

Olshanskii, M. et al. (2009) ‘Grad-div stabilization and subgrid pressure models for the incompressible Navier-Stokes equations’, Computer Methods in Applied Mechanics and Engineering, 198(49-52), pp. 3975–3988. doi: 10.1016/j.cma.2009.09.005.

Raghunandan, A. et al. (2021) ‘Bulk flow of cerebrospinal fluid observed in periarterial spaces is not an artifact of injection’, eLife, 10, pp. 1–15. doi: 10.7554/eLife.65958.

Rajagopal, K. R., Wineman, A. S. and Gandhi, M. (1986) ‘On boundary conditions for a certain class of problems in mixture theory’, International Journal of Engineering Science, 24(8), pp. 1453–1463. doi: 10.1016/0020-7225(86)90074-1.

Rasmussen, M. K., Mestre, H. and Nedergaard, M. (2021) ‘Fluid Transport in the Brain’, Physiological Reviews.

Schain, A. J. et al. (2017) ‘Cortical spreading depression closes the paravascular space and impairs glymphatic flow: Implications for migraine headache’, The Journal of Neuroscience, 37(11), pp. 3390–16. doi: 10.1523/JNEUROSCI.3390-16.2017.

Schley, D. et al. (2006) ‘Mechanisms to explain the reverse perivascular transport of solutes out of the brain’, Journal of Theoretical Biology, 238(4), pp. 962–974. doi: 10.1016/j.jtbi.2005.07.005.

Selkoe, D. J. and Hardy, J. (2016) ‘The amyloid hypothesis of Alzheimer’s disease at 25 years’, EMBO Molecular Medicine, 8(6), pp. 595–608. doi: 10.15252/emmm.201606210.

Senjuntichai, T. and Rajapakse, R. K. N. D. (1993) ‘Transient response of a circular cavity in a poroelastic medium’, International Journal for Numerical and Analytical Methods in Geomechanics, 17(6), pp. 357–383. doi: 10.1002/nag.1610170602.

Shih, A. Y. et al. (2012) ‘Two-photon microscopy as a tool to study blood flow and neurovascular coupling in the rodent brain’, Journal of Cerebral Blood Flow and Metabolism, 32(7), pp. 1277–1309. doi: 10.1038/jcbfm.2011.196.

Shih, A. Y. et al. (2015) ‘Robust and fragile aspects of cortical blood flow in relation to the underlying angioarchitecture’, Microcirculation, 22(3), pp. 204–218.

Shim, J. J. et al. (2021) ‘Finite Element Implementation of Biphasic-Fluid Structure Interactions in febio’, Journal of biomechanical engineering, 143(9). doi: 10.1115/1.4050646.

Shim, J. J. and Ateshian, G. A. (2021) ‘A hybrid biphasic mixture formulation for modeling dynamics in porous deformable biological tissues’, Archive of Applied Mechanics, vi, pp. 13–16. doi: 10.1007/s00419-020-01851-8.

Silva, A. C., Koretsky, A. P. and Duyn, J. H. (2007) ‘Functional MRI impulse response for BOLD and CBV contrast in rat somatosensory cortex’, Magnetic Resonance in Medicine: An Official Journal of the International Society for Magnetic Resonance in Medicine, 57(6), pp. 1110–1118.

Smith, A. J. et al. (2017) ‘Test of the ‘glymphatic’ hypothesis demonstrates diffusive and aquaporin-4-independent solute transport in rodent brain parenchyma’, eLife, 6, pp. 1–16. doi: 10.7554/eLife.27679.

Smith, A. J. and Verkman, A. S. (2019) ‘CrossTalk opposing view: Going against the flow: interstitial solute transport in brain is diffusive and aquaporin-4 independent’, The Journal of Physiology, 0, pp. 1–4. doi: 10.1113/jp277636.

Smith, J. H. and Humphrey, J. A. C. (2007) ‘Interstitial transport and transvascular fluid exchange during infusion into brain and tumor tissue’, Microvascular Research, 73(1), pp. 58–73. doi: 10.1016/j.mvr.2006.07.001.

Støverud, K. H. et al. (2013) ‘CSF pressure and velocity in obstructions of the subarachnoid spaces’, The neuroradiology journal, 26(2), pp. 218–226.

Sykova, E. et al. (2008) ‘Diffusion in Brain Extracellular Space’, Physiol Rev., 88(4), pp. 1277–1340. doi: 10.1152/physrev.00027.2007.

Thomas, J. H. (2019) ‘Fluid dynamics of cerebrospinal fluid flow in perivascular spaces’, Journal of the Royal Society Interface, 16(159). doi: 10.1098/rsif.2019.0572.

Tithof, J. et al. (2019) ‘Hydraulic resistance of periarterial spaces in the brain’, Fluids and Barriers of the CNS, 16(1), pp. 1–13. doi: 10.1186/s12987-019-0140-y.

Treloar, L. R. G. (1975) The physics of rubber elasticity. Oxford University Press, USA.

Troyetsky, D. E. et al. (2021) ‘Dispersion as a waste-clearance mechanism in flow through penetrating perivascular spaces in the brain’, Scientific Reports, 11(1), pp. 1–12. doi: 10.1038/s41598-021-83951-1.

Tseng, B. P. et al. (1999) ‘Deposition of monomeric, not oligomeric, Aβ mediates growth of Alzheimer’s disease amyloid plaques in human brain preparations’, Biochemistry, 38(32), pp. 10424–10431. doi: 10.1021/bi990718v.

Turner, K. L. et al. (2020a) ‘Neurovascular coupling and bilateral connectivity during NREM and REM sleep’, bioRxiv.

Turner, K. L. et al. (2020b) ‘Neurovascular coupling and bilateral connectivity during NREM and REM sleep’, Elife, 9, p. e62071.

van Veluw, S. J. et al. (2020) ‘Vasomotion as a Driving Force for Paravascular Clearance in the Awake Mouse Brain’, Neuron, 105(3), pp. 549–561.e5. doi: 10.1016/j.neuron.2019.10.033.

Vinje, V., Bakker, E. N. T. P. and Rognes, M. E. (2021) ‘Brain solute transport is more rapid in periarterial than perivenous spaces’, Scientific Reports, (0123456789), pp. 1–11. doi: 10.1038/s41598-021-95306-x.

Wang, P. and Olbricht, W. L. (2011) ‘Fluid mechanics in the perivascular space’, Journal of Theoretical Biology, 274(1), pp. 52–57. doi: 10.1016/j.jtbi.2011.01.014.

Weickenmeier, J. et al. (2018) ‘Brain stiffens post mortem’, Journal of the Mechanical Behavior of Biomedical Materials, 84(January), pp. 88–98. doi: 10.1016/j.jmbbm.2018.04.009.

Weller, R. O. (1998) ‘Pathology of cerebrospinal fluid and interstitial fluid of the CNS: significance for Alzheimer disease, prion disorders and multiple sclerosis’, Journal of neuropathology and experimental neurology, 57(10), pp. 885–894.

Weller, R. O., Kida, S. and Zhang, E. -T (1992) ‘Pathways of Fluid Drainage from the Brain - Morphological Aspects and Immunological Significance in Rat and Man’, Brain Pathology, 2(4), pp. 277–284. doi: 10.1111/j.1750-3639.1992.tb00704.x.

Winder, A. T. et al. (2017) ‘Weak correlations between hemodynamic signals and ongoing neural activity during the resting state’, Nature Neuroscience, 20(12), pp. 1761–1769. doi: 10.1038/s41593-017-0007-y.

Xie, K. H., Liu, G. Bin and Shi, Z. Y. (2004) ‘Dynamic response of partially sealed circular tunnel in viscoelastic saturated soil’, Soil Dynamics and Earthquake Engineering, 24(12), pp. 1003–1011. doi: 10.1016/j.soildyn.2004.05.005.

Xie, L. et al. (2013) ‘Sleep drives metabolite clearance from the adult brain.’, Science, 342(6156), pp. 373–377. doi: 10.1126/science.1241224.

Yamada, M. (2000) ‘Cerebral amyloid angiopathy: An overview’, Neuropathology, 20(1), pp. 8–22. doi: 10.1046/j.1440-1789.2000.00268.x.

Yamada, M. (2015) ‘Cerebral amyloid angiopathy: Emerging concepts’, Journal of Stroke, 17(1), pp. 17–30. doi: 10.5853/jos.2015.17.1.17.

Yetkin, F. et al. (2010) ‘Cerebrospinal fluid viscosity: a novel diagnostic measure for acute meningitis’, South Med J, 103(9), pp. 892–895.

Yu, X. et al. (2014) ‘Deciphering laminar-specific neural inputs with line-scanning fMRI’, Nature Methods, 11(1), pp. 55–58. doi: 10.1038/nmeth.2730.

Yu, X. et al. (2016) ‘Sensory and optogenetically driven single-vessel fMRI’, Nature Methods, 13(4), pp. 337–340. doi: 10.1038/nmeth.3765.

Zhang, Q. et al. (2019) ‘Cerebral oxygenation during locomotion is modulated by respiration’, Nature Communications, 10(1). doi: 10.1038/s41467-019-13523-5.

