## Supplemental figures and captions for "Arterial vasodilation drives convective fluid flow in the brain: a poroelastic model"

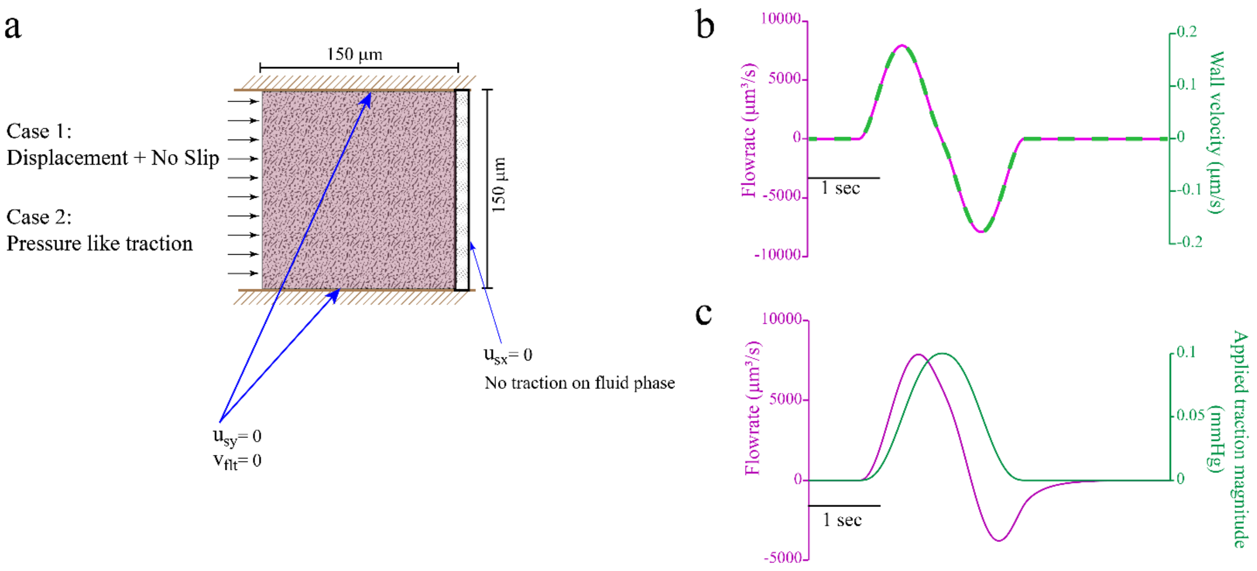


Fig. S1: 2D poroelastic model demonstrates the difference between SAS and ECS fluid exchange during arteriolar dilation.

**a.** 2D Poroelastic model dimensions and boundary conditions. All other parameters were the same as the ones used for the brain tissue in the rest of the article. **b.** Flow rate through the right edge has the same waveform as the wall velocity of the left edge for Case 1, where a displacement boundary condition was imposed on the left edge along with a no-slip condition. **c.** Flow rate through the right edge lags the changes in the pressure-like traction at the left edge for Case 2, where a pressure-like traction was applied on the left edge.

Case 1, where a direct displacement boundary condition was imposed represents flow through the SAS during arteriolar dilation. Case 2 represents flow through the ECS, where fluid flow is induced by pressure changes in the PVS.

Note: The flow rate was calculated by assuming a $150\mu m$ thickness perpendicular to the plane.

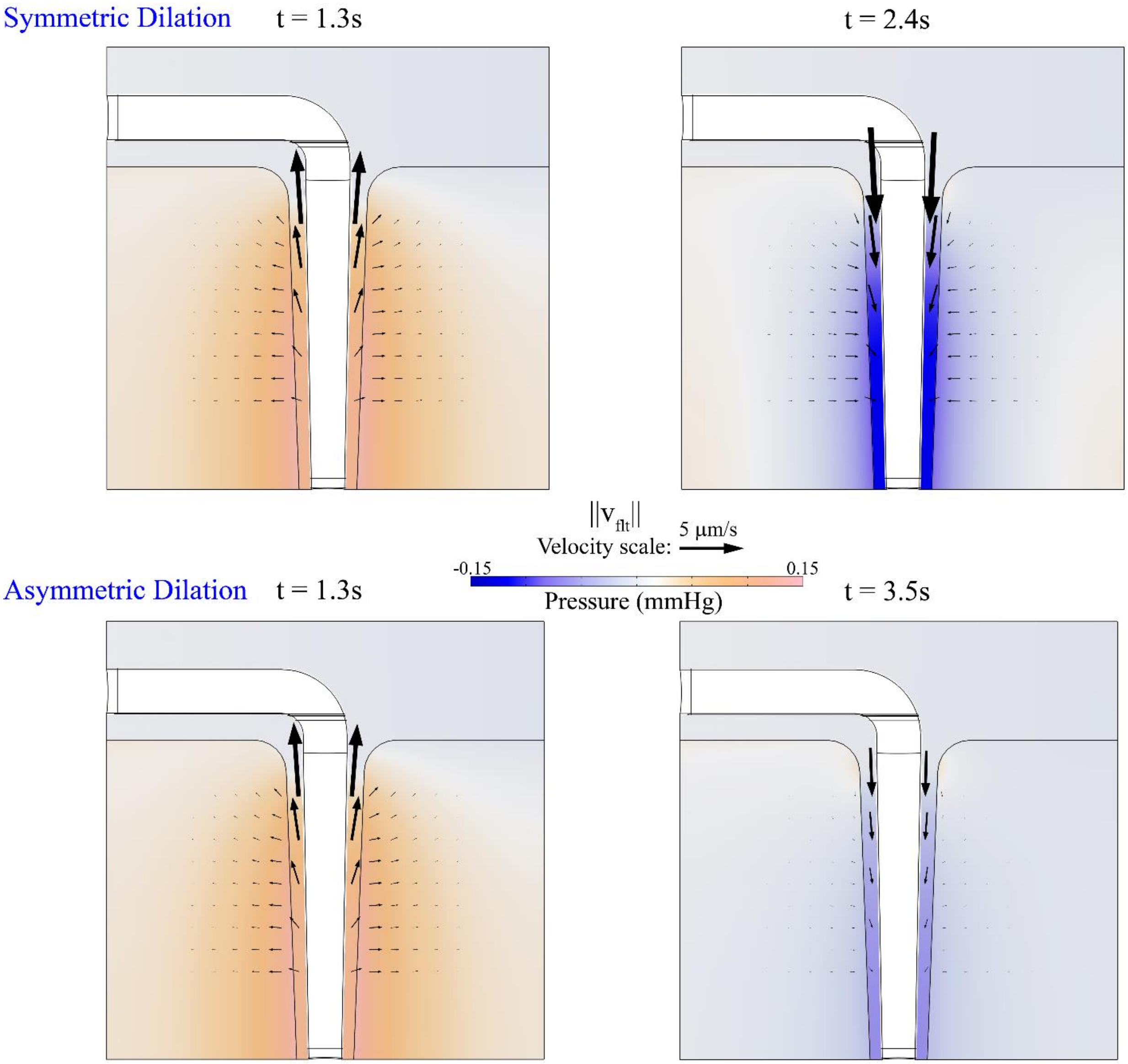


Fig. S2: Filtration velocity for temporally symmetric and asymmetric dilation

The pressure and filtration velocity in the PVS and the ECS at the times of maximum radially outward and inward arteriolar wall velocity for symmetric (top) and asymmetric (bottom) dilation. The colors show the pressure value in mmHg and the arrows show the magnitude and direction of the filtration of the filtration velocity. Fewer rows of arrows were used in the PVS to avoid overlapping arrows. The continuity of filtration velocity across the two regions is best demonstrated by the bottom most row, where the fluid flows more in the radial direction than the axial direction.

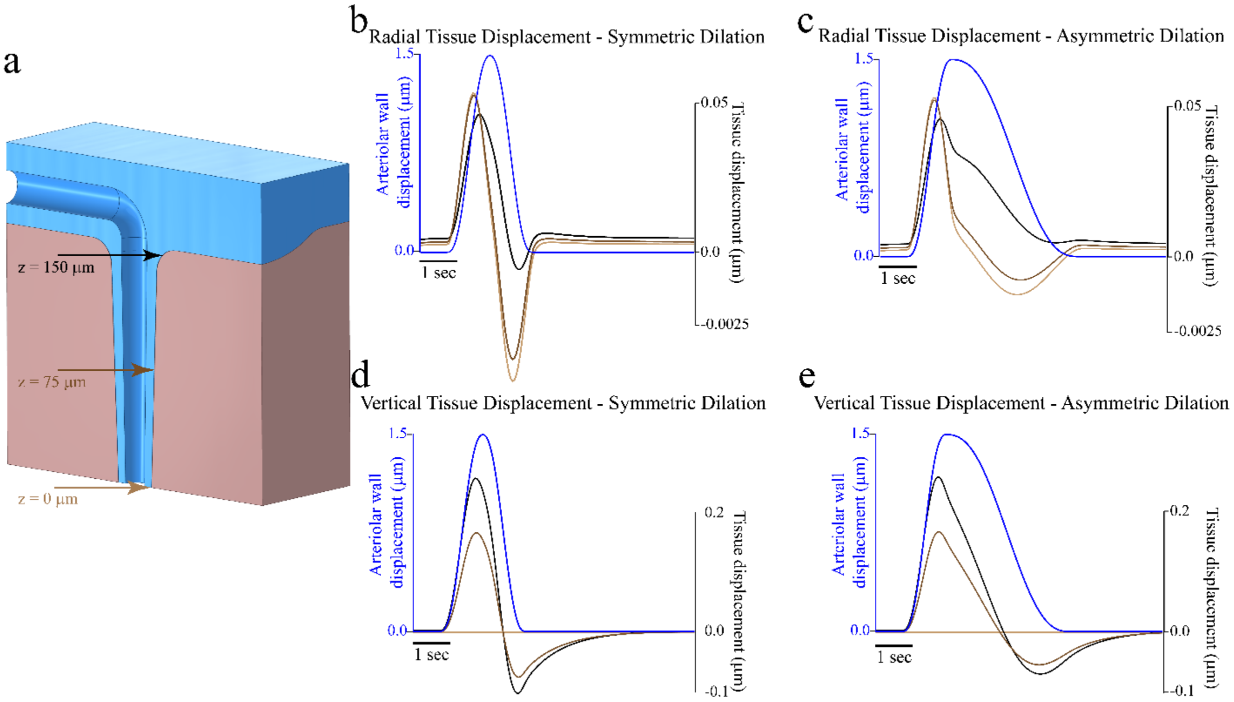


Fig. S3: Dilation of the brain tissue in the model at the PVS-ECS interface in the radial direction **b-c** and in the vertical direction **e-f.** The three locations where the displacement was calculated is shown in **a.** The blue line in all the subplots is the arteriolar wall dilation in the radial direction.

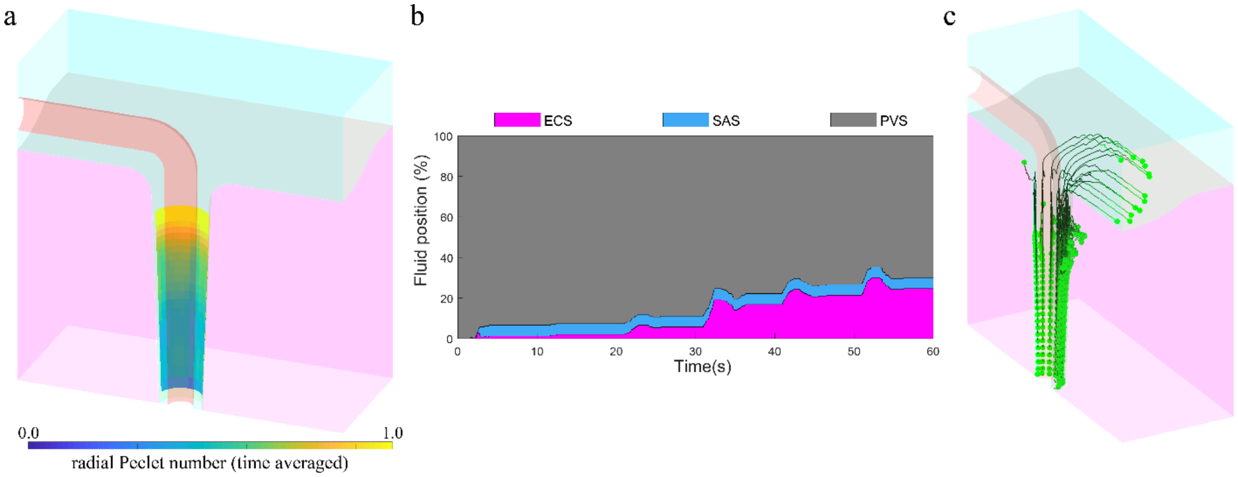


Fig. S4: Directional fluid flow from the PVS into the ECS driven by vasodilation is not an artifact of the imposed pressure difference across the SAS

The simulations with the asymmetric vasodilation (bottom part of Fig. 3) were repeated with a smaller pressure difference ($p_0 = 0.001\, mmHg$) across the SAS. The radial Peclet number averaged over 10s of simulation with a single 5-second-long vasodilation event **(a)** show that directional fluid flow driven by the temporally asymmetric waveform of functional hyperemia is not an artifact of the fluid flow in the SAS. This is confirmed by examining the PVS fluid position **(b)** and trajectories **(c)** in a particle tracking simulation performed for 60s, where a single vasodilation event is repeated once every 10s.

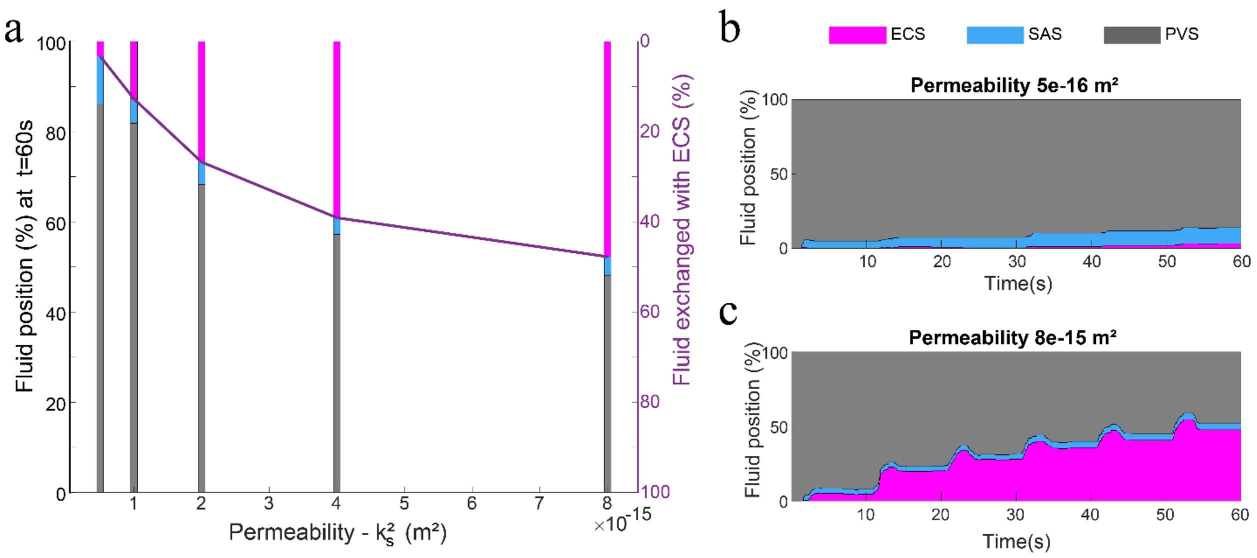


Fig. S5: PVS fluid penetration into the ECS increases with increased brain fluid permeability ($k_s^2$).

**a.** PVS Fluid distribution at the end of a 60 second fluid particle tracking simulation. The fluid exchanged with ECS (magenta) increases with increased permeability, while fluid exchanged with SAS (blue) decreases. **b.** and **c.** Show the fluid distribution during the 60 seconds for $k_s^2 = 0.5\ \times 10^{-15} m^2$ and $k_s^2 = 8.0\ \times 10^{-15} m^2$ respectively.

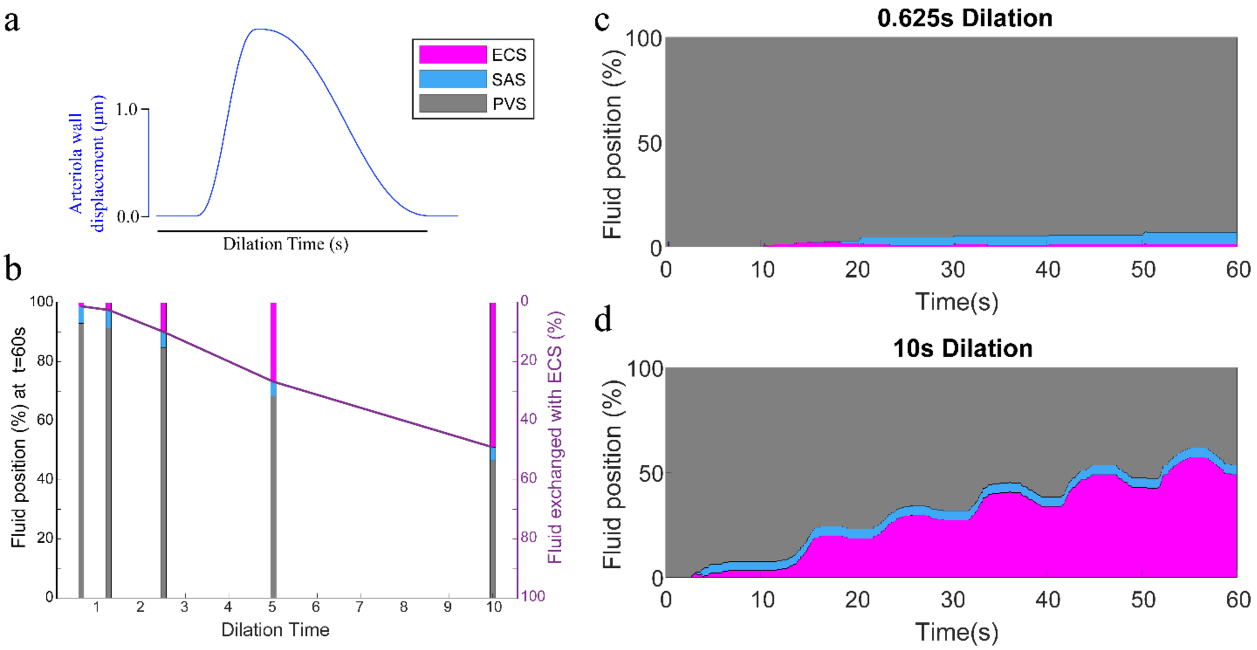


Fig. S6: PVS fluid penetration into the ECS is higher for low frequency vasodilation.

a. The arteriolar dilation waveform showing the dilation time. One dilation event was used for 10 seconds of simulation. b. PVS Fluid distribution at the end of a 60 second fluid particle tracking simulation. The fluid exchanged with ECS (magenta) is higher for slower dilation, while fluid exchanged with SAS (blue) does not change appreciably with dilation frequency. **c.** and **d.** Show the fluid distribution during the 60 seconds for dilation time of 0.625s and 10s respectively.

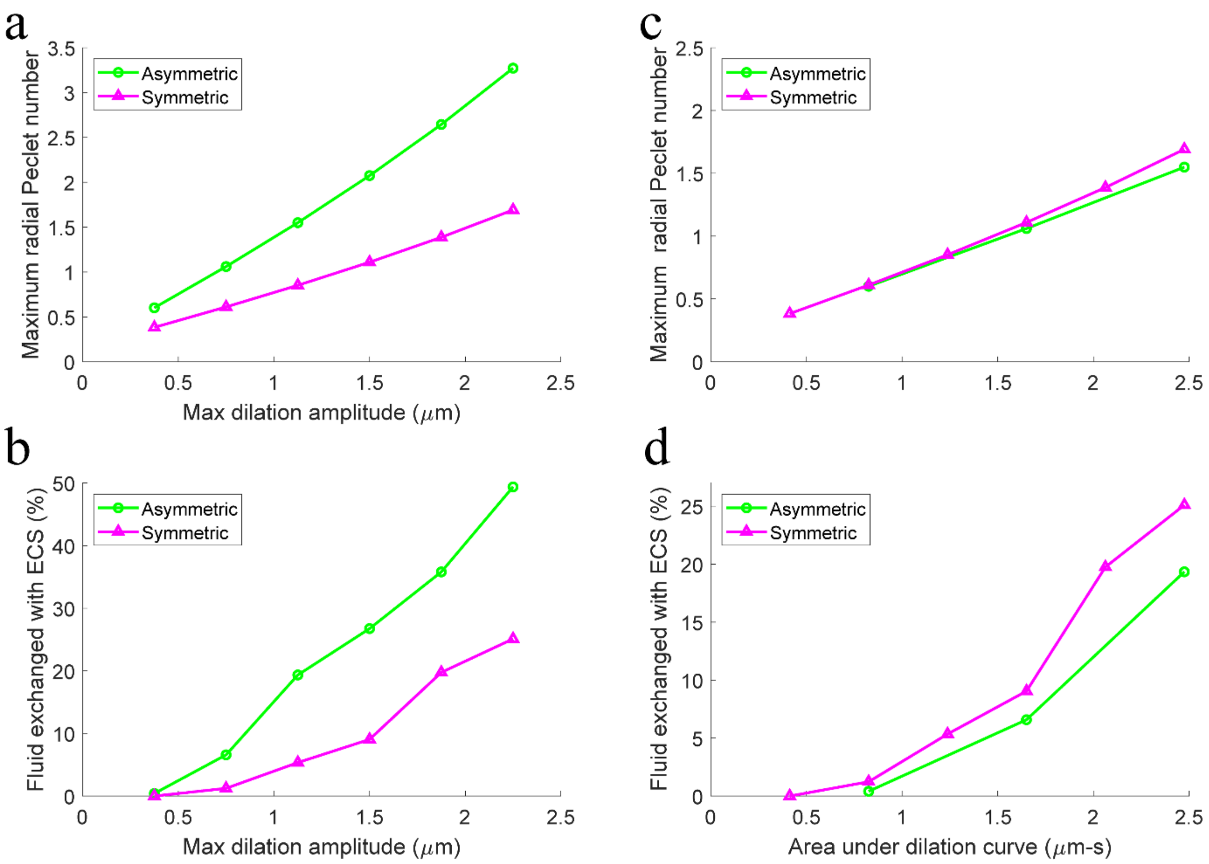


Fig S7: The area under the dilation curve, not the maximum dilation amplitude, is an indicator of directional PVS fluid flows into the ECS.

**a.** and **b.** show that the maximum time-averaged radial Peclet number ($Pe_r$) and PVS fluid exchanged with the ECS (over 60 seconds with one dilation per 10 seconds), which are measures of directional PVS fluid flow into the ECS, are appreciably affected by the on the waveform of the vasodilation for the same peak dilation value.

**c.** and **d.** show that the directional PVS fluid flow into the ECS is not appreciably affected by the waveform, when the area under the dilation curve is kept the same.
